# Cohesin occupancy and composition at enhancers and promoters are linked to DNA replication origin proximity in *Drosophila*

**DOI:** 10.1101/539270

**Authors:** Michelle Pherson,, Ziva Misulovin, Maria Gause, Dale Dorsett

## Abstract

Cohesin consists of the Smc1-Smc3-Rad21 tripartite ring and the SA protein that interacts with Rad21. The Nipped-B protein loads cohesin topologically around chromosomes to mediate sister chromatid cohesion and facilitate long-range control of gene transcription. It is largely unknown how Nipped-B and cohesin associate specifically with gene promoters and transcriptional enhancers, or how sister chromatid cohesion is established. Here we use genome-wide chromatin immunoprecipitation in *Drosophila* cells to show that SA and the Fs(1)h (BRD4) BET domain protein help recruit Nipped-B and cohesin to enhancers and DNA replication origins, while the MED30 subunit of the Mediator complex directs Nipped-B and Rad21 to promoters. All enhancers and their neighboring promoters are close to DNA replication origins and bind SA with proportional levels of cohesin subunits. Most promoters are far from origins and lack SA, but bind Nipped-B and Rad21 with sub-proportional amounts of Smc1, indicating that they bind SA-deficient cohesin part of the time. Genetic data confirm that Nipped-B and Rad21 function together with Fs(1)h in vivo to facilitate *Drosophila* development. These findings demonstrate that Nipped-B and cohesin are differentially targeted to enhancers and promoters and suggest models for how SA and DNA replication help establish sister chromatid cohesion and facilitate enhancer-promoter communication. They indicate that SA is not an obligatory cohesin subunit but a factor that controls cohesin location on chromosomes.

## Introduction

Cohesin mediates sister chromatid cohesion to ensure accurate chromosome segregation and also plays roles in DNA repair and gene transcription (Dorsett and Ström 2012; Dorsett and Merkenschlager 2013; Uhlmann 2016, Morales and Losada 2018; Villa‐Hernández and Bermejo 2018). In *Drosophila*, cohesin facilitates enhancer-promoter communication and regulates activity of the Polycomb repressive complex 1 at silenced and active genes (Rollins et al. 1999; Schaaf et al. 2013a; Schaaf et al. 2013b; Pherson et al. 2017).

Cohesin structure and chromosome binding are relatively well-understood. The Smc1, Smc3 and Rad21 subunits form a tripartite ring and SA interacts with Rad21. A Nipped-B - Mau2 complex loads cohesin topologically around chromosomes and a Pds5 - Wapl complex removes cohesin. SA, Nipped-B, Pds5 and Wapl contain HEAT repeats and interact with cohesin to control its binding and activities (Neuwald and Hirano 2000; Wells et al. 2017). These accessory proteins facilitate ring opening to load and remove cohesin from chromosomes (Murayama and Uhlmann 2014; Çamdere et al. 2015; Murayama and Uhlmann 2015; Beckouët et al. 2016; Elbatsh et al. 2016; Yu 2016, Ouyang and Yu 2017; Petela et al. 2018).

Less is known about how cohesin is targeted to sequences that control gene transcription or how sister chromatid cohesion is established. In *Drosophila*, cohesin associates with active genes, transcriptional enhancers, and the Polycomb response elements (PREs) that control epigenetic gene silencing (Misulovin et al. 2008; Schaaf et al. 2013a; Schaaf et al. 2013b; Swain et al. 2016; Misulovin et al. 2018). Cohesin occupies all enhancers and PREs, and preferentially those active genes positioned within several kilobases of the early DNA replication origins (MacAlpine et al. 2010; Misulovin et al. 2018).

The Pds5 and Wapl cohesin removal factors limit the size of cohesin domains surrounding early origins, while Pds5 and the Brca2 DNA repair protein, which form a complex lacking Wapl (Brough et al. 2012; Kusch 2015) have opposing effects on SA origin occupancy and sister chromatid cohesion (Misulovin et al. 2018). Pds5 is required for sister chromatid cohesion and facilitates SA binding, while Brca2 inhibits SA binding and counters the ability of Pds5 to support sister cohesion when Pds5 levels are low. These findings gave rise to the idea that Pds5 and SA function at replication origins to establish chromatid cohesion (Misulovin et al. 2018).

To gain more insight into how cohesin associates with gene regulatory sequences we used genome-wide chromatin immunoprecipitation (ChIP-seq) to investigate how multiple cohesin subunits occupy different genomic features in *Drosophila* cells. We also examined the roles of cohesin subunits, the Mediator complex, and the Fs(1)h (BRD4) BET domain protein in cohesin localization. The results indicate that cohesin associates with enhancers and most promoters by different mechanisms and that proximity to DNA replication origins influences cohesin occupancy and composition.

## Results

We compared how cohesin subunits and the Nipped-B cohesin loading factor (Fig. 1A) occupy promoters, enhancers, and Polycomb Response Elements (PREs) by ChIP-seq in ML-DmBG3-c2 (BG3) cells derived from 3^rd^ instar central nervous system. Multiple biological replicates were used for each protein (Supplemental Table S1). Fig. 1B shows ChIP-seq in a region near an early DNA replication origin where cohesin levels are high and Fig. 1C shows an origin-distal region with lower occupancy. Preimmune serum ChIP-seq shows insignificant enrichment of functional features (Fig. 1, Supplemental Fig. S1).

**Figure 1.**
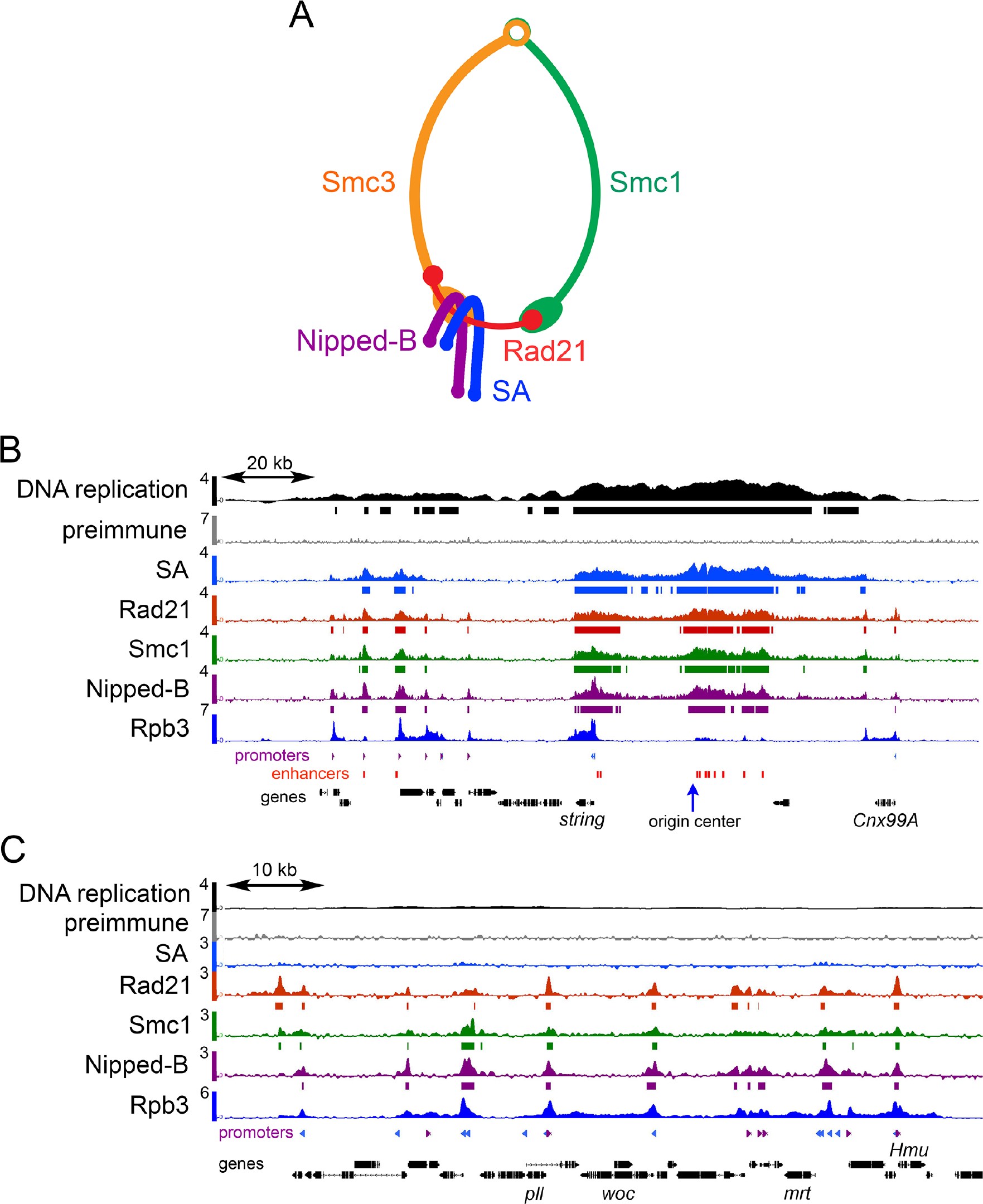
Cohesin and Nipped-B ChIP-seq in BG3 cells. (*A*) Cohesin subunit structure. (*B*) ChIP-seq near an early DNA replication origin at the *string* (*cdc25*) gene. ChIP-seq and DNA replication data are plotted as log_2_ enrichment. Bars under each track indicate enrichment in the 95^th^ percentile over regions ≥300 bp. Rpb3 RNA polymerase II subunit data is from a prior publication (Pherson et al. 2017). Locations of promoters (purple and blue, forward and reverse) and enhancers (red) are indicated underneath the tracks. (*C*) ChIP-seq in a region distant from an early replication origin containing the *woc* (*without children*) gene.

### Cohesin subunits occupy functional features in different proportions

Fig. 2A illustrates the distributions of the SA, Rad21, Smc1 and Nipped-B occupancies of promoters (PRO) enhancers (ENH) PREs (PRE) and centers of early DNA replication origins (ORI) using violin plots. All show insignificant median enrichment across 6,892 randomly positioned sequences (RAN). SA does not occupy most of the 7,389 active promoters, but essentially all 2,353 enhancers, 195 PREs, and 78 origin centers. In contrast, Rad21, Smc1 and Nipped-B occupy most active promoters and all enhancers, PREs and origins. SA has the highest median occupancy at origins while the Rad21 and Smc1 ring components are highest at enhancers and Nipped-B is maximal at PREs (Fig. 2A, Supplemental Table S2).

**Figure 2.**
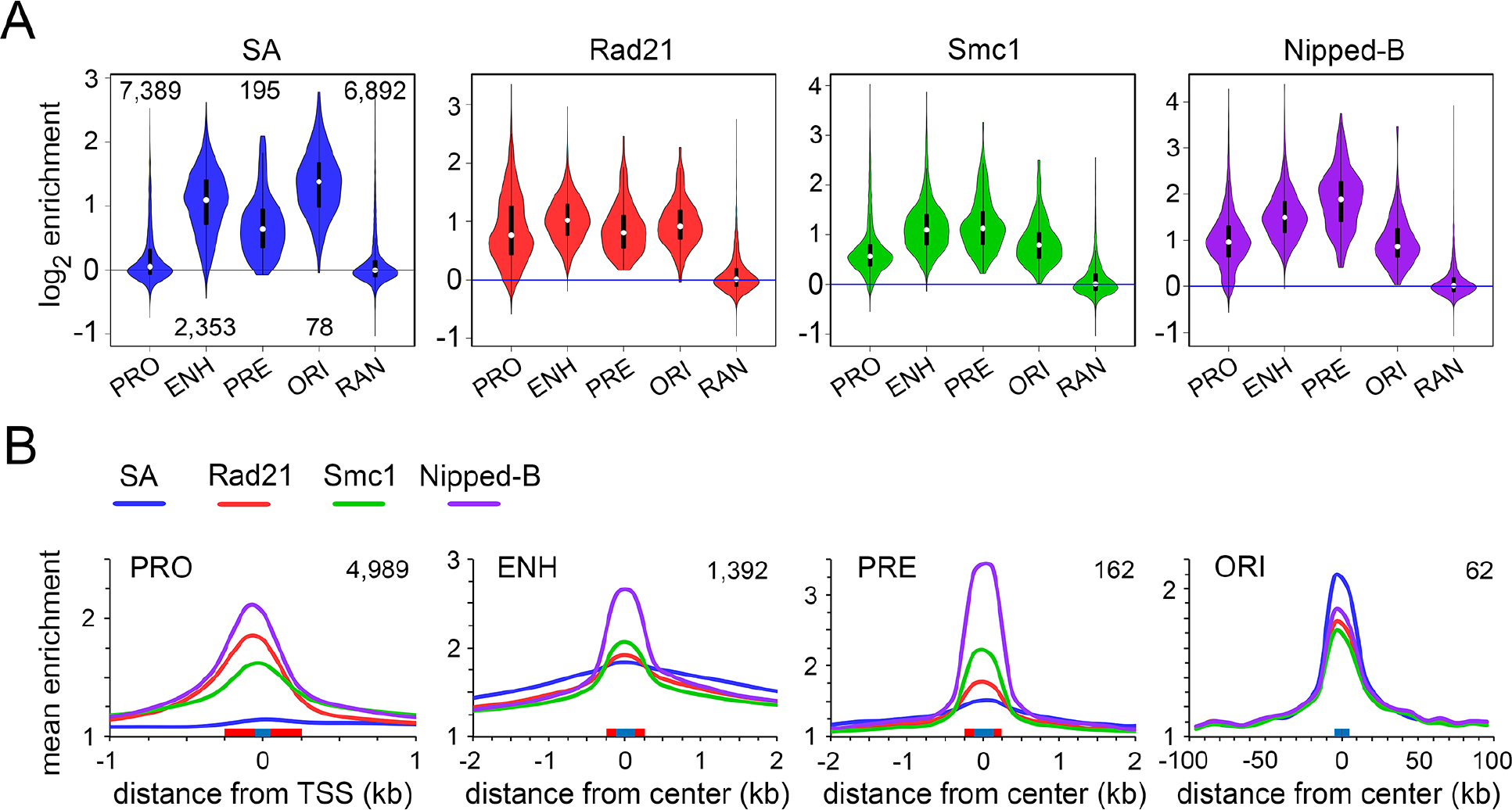
Cohesin subunits are present in different ratios at promoters, enhancers, PREs and replication origins in BG3 cells. (*A*) Violin plot distributions of SA (blue) Rad21 (red) Smc1 (green) and Nipped-B (purple) at active promoters (PRO) enhancers (ENH) Polycomb Response Elements (PRE) early replication origin centers (ORI) and randomly positioned sequences (RAN). The numbers of each type of feature analyzed are indicated in the SA plot. White dots are the median values given in Supplemental Table S2. (*B*) Meta-analyses of promoters, enhancers, PREs and early replication origins for SA (blue) Rad21 (red) Smc1 (green) and Nipped-B (purple). Red boxes on the X axes indicate the feature sizes used to calculate occupancy for the violin plots and blue boxes indicate the bin sizes used to average enrichment for the meta-analysis. The numbers of each type of feature used for meta-analysis are indicated in the upper right corner of each graph. These are less than for the violin plots because features that overlap in the meta-analysis region were removed to minimize distortions.

Differential occupancy by cohesin subunits is also illustrated by meta-analyses that average the distribution of ChIP-seq enrichment centered at each type of feature (Fig. 2B). Of the four proteins, Nipped-B (purple) shows the highest mean enrichment at promoters, enhancers, PREs, and the second highest at origins (Fig. 2B). SA is the highest at origins. At promoters, there is minimal SA (blue) and Rad21 enrichment (red) is higher than Smc1 (green). Rad21 enrichment is also higher than Smc1 at origin centers. In contrast, Smc1 shows higher enrichment than Rad21 at enhancers and PREs. At enhancers, SA extends into the flanking regions more than Rad21 and Smc1, suggesting that some SA binds independently of cohesin (Fig. 2B).

Enrichment values for the individual cohesin subunits depend on different efficiencies of crosslinking and precipitation and thus cannot be directly compared. However, we infer that the stoichiometry of the subunits varies between the different features because their relative mean enrichments differ. As described below, depletion experiments confirm that epitope masking or differential crosslinking are not responsible for low SA and Smc1 levels seen at most promoters.

SA shows higher mean enrichment relative to the other cohesin subunits in meta-origin analysis but lower enrichment at promoters and enhancers (Fig. 2B). This indicates that cohesin at features close to origins has more SA than cohesin at origin-distal features and/or that some SA binds near origins independently of cohesin. As described below, we find that enhancers and the rare promoters that bind SA are origin-proximal and that promoters with low SA are origin-distal.

### Nipped-B and Rad21 occupy most gene promoters without SA

Fig. 1C shows origin-distal promoters occupied by Nipped-B, Rad21 and Smc1 with little or no SA. Dot plots of Rad21 promoter occupancy against SA (Fig. 3A) or Smc1 (Fig. 3B) show that most promoters have Rad21, sub-proportional Smc1, and no SA (black arrows). We defined “high SA” promoters as those within regions of SA enrichment in the 95^th^ percentile. These regions, ranging from 300 bp to several kilobases, are marked by bars underneath the SA ChIP-seq track in Fig. 1B. High SA promoters represent 12% of active promoters (895 / 7,398) and are plotted in red in the dot plots (Fig. 3A,B). In contrast to most promoters, high SA promoters have proportional SA, Rad21 and Smc1 levels similar to enhancers and PREs (Fig. 3A,B). This implies that cohesin subunits are more stoichiometric at high SA promoters and enhancers than at most promoters.

**Figure 3.**
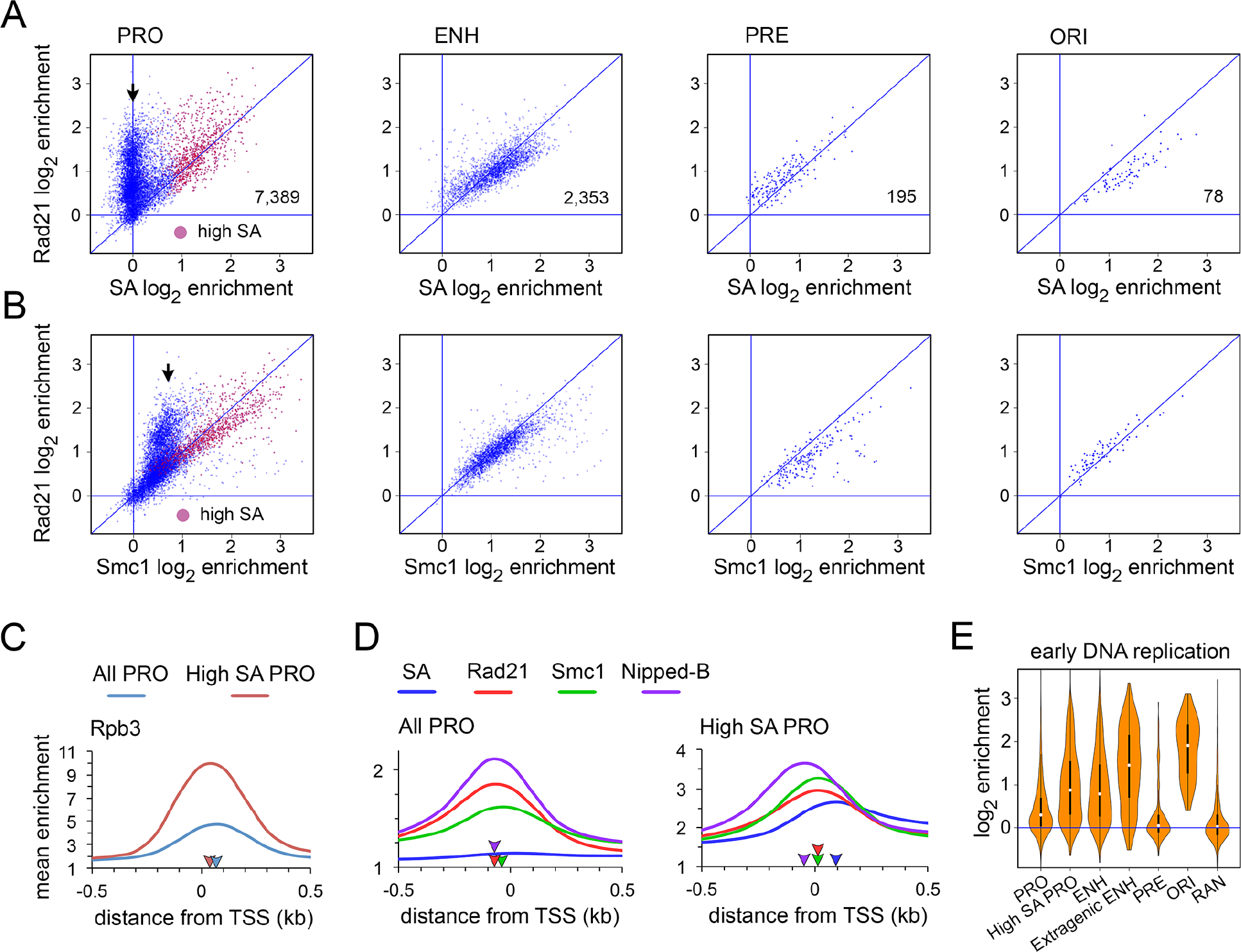
Nipped-B, Rad21 and sub-proportional Smc1 occupy most active promoters without SA in BG3 cells. (*A*) Dot plots of SA versus Rad21 enrichment at promoters (PRO) enhancers (ENH) PREs (PRE) and origin centers (ORI). The numbers of each feature type are indicated in the plots. Promoters with high SA enrichment (95^th^ percentile over regions ≥300 bp) are plotted in red. The black arrow indicates promoters with Rad21 but no SA. (*B*) Dot plots of Smc1 enrichment vs. Rad21 enrichment at the indicated feature types. The black arrow indicates promoters with sub-proportional Smc1. (*C*) Rpb3 (Pol II) promoter meta-analysis for active promoters (blue) and the subset occupied by SA (high SA, red). Blue and red arrowheads on the X-axis indicate the positions of peak enrichment. (*D*) SA (blue) Rad21 (red) Smc1 (green) and Nipped-B (purple) promoter meta-analysis for all promoters (All PRO) and the subset occupied by SA (High SA PRO). The purple, red, green, and blue arrowheads on the X-axes indicate peak enrichment for Nipped-B, Rad21, Smc1 and SA. SA peak enrichment is indicated only for high SA promoters. (*E*) Violin plots showing enrichment of early-replicating DNA for all active promoters (PRO) promoters occupied by SA (High SA PRO) enhancers (ENH) the subset of enhancers positioned at least 500 bp outside of a transcribed region (Extragenic ENH) PREs (PRE) centers of replication origins (ORI) and randomly positioned sequences (RAN).

Sub-proportional Smc1 levels at SA-deficient promoters implies that Nipped-B and Rad21 occupy these promoters in both the absence and presence of Smc1-Smc3 dimers. We envision that a percentage of each of these promoters in a cell population bind Nipped-B - Rad21 complexes without Smc1 and Smc3, while another fraction has Smc1-Smc3-Rad21 tripartite rings. In contrast, enhancers and high SA promoters are primarily occupied by cohesin complexes containing SA.

Meta-analysis using Rpb3 ChIP-seq data (Pherson et al. 2017) shows that high SA promoters (red) have more RNA polymerase on average than most promoters (blue) (Fig. 3C). Rpb3 peaks downstream of the transcription start site at +30 bp for high SA promoters and at +65 for all promoters (red and blue arrowheads). This agrees with PRO-seq studies showing that genes with more cohesin show above average transcription and promoter-proximal Pol II pausing (Schaaf et al. 2013a).

Unexpectedly, the positions of different cohesin subunits relative to the transcription start site differ at promoters. Averaging all promoters, Nipped-B and Rad21 peak 70 bp upstream of the start site (−70 bp, purple and red arrowheads) while Smc1 peaks at −35 bp (green arrowhead) (Fig. 3D). We interpret this as indicating that some Rad21 binds to promoters independently and upstream of Smc1 and that another fraction interacts with Smc1 downstream in Smc1-Smc3-Rad21 cohesin complexes. The close alignment of the Nipped-B and Rad21 peaks supports the idea that these promoters are occupied by Nipped-B – Rad21 complexes.

Cohesin is positioned further downstream at high SA promoters compared to most promoters (Fig. 3D). Nipped-B peaks upstream at −50 bp (purple arrowhead) but Rad21 (red) and Smc1 (green) peak together just downstream of the transcription start site, and SA (blue) peaks further downstream near +100 (Fig. 3D). SA also extends more than Rad21 and Smc1 into the transcribed region. Pol II (Rpb3) peaks 30 bp downstream of Rad21 and Smc1 but nearly 70 bp upstream of SA. We theorize that some SA associates with the elongating Pol II complex that enters into the gene body. The precise alignment of Rad21 and Smc1 peaks at high SA promoters contrasts with their misalignment at most promoters and correlates with their more proportional levels. We interpret this as indicating that high SA promoters are occupied primarily by complete cohesin complexes.

### High SA promoters and enhancers are close to early DNA replication origins

As noted above, SA levels are highest near early replication origins suggesting that high SA promoters are positioned close to origins and those with low SA are located farther away. We tested this by comparing the levels of early S phase DNA synthesis at functional features. Those close to early origins should be replicated in early S phase. We measured early DNA synthesis by incorporation of the EdU thymidine analog and high-throughput sequencing in cells blocked in early S phase with hydroxyurea. The results are similar to data obtained using BrdU and microarrays (Eaton et al. 2011). As predicted, high SA promoters and enhancers experience higher DNA replication in early S phase than most promoters (Fig. 3E). PREs and randomly-positioned sequences show low replication while origin centers show high levels, as expected.

Extragenic enhancers located outside of transcribed regions experience higher early S phase DNA synthesis than most enhancers, indicating that they are positioned particularly close to early origins (Fig. 3E). The origin recognition complex (ORC) recruits the MCM2-7 helicase complex that unwinds duplex DNA to start replication during origin licensing in early G1 and transcription pushes MCM2-7 to regions outside of genes (Powell et al. 2015). Repositioning of MCM2-7 outside of transcribed regions can explain higher early DNA synthesis at extragenic enhancers and raises the possibility that enhancers may help position MCM2-7. Supplemental Fig. S2A illustrates striking overlaps of early origins with clusters of enhancers.

### SA facilitates cohesin and Nipped-B occupancy of enhancers and origin-proximal promoters

We depleted SA by RNAi in BG3 cells and conducted ChIP-seq for Nipped-B, Rad21 and Smc1 to test if SA positions cohesin at origin-proximal features. RNAi was conducted for 3 to 4 days, depleting SA by 80 to 90% without reducing sister chromatid cohesion and slightly slowing cell proliferation (Schaaf et al. 2009; Misulovin et al. 2018) (Supplemental Fig. S3A). SA depletion partially reduces total Rad21 protein and does not alter total Nipped-B levels (Schaaf et al. 2009) (Supplemental Fig. S3A). BG3 cells divide during the depletion with a 24 hour cycle and changes revealed by ChIP-seq reflect an altered steady-state equilibrium of cohesin occupancy.

Fig. 4 shows that SA depletion shifts cohesin occupancy from origin-proximal to origin-distal features supporting the idea that SA positions cohesin close to origins. Rad21 (Fig. 4A) Smc1 (Fig. 4B) and Nipped-B (Fig. 4C) increase at most promoters upon SA depletion and decrease at enhancers and origin-proximal regions. They also decrease at PREs with the exception that Smc1 PRE occupancy is only slightly modified. The changes of cohesin occupancy are statistically significant except for Smc1 at PREs (Supplemental Table S2). Supplemental Fig. S2B illustrates how Rad21, Smc1 and Nipped-B decrease at the *string* enhancers and flanking promoters near a replication origin. In contrast, Rad21 and Smc1 increase slightly at some origin-distal promoters in Supplemental Fig. S2C. Promoter dot plots in Fig. 4 show that although Rad21, Smc1 and Nipped-B decrease at the origin-proximal high SA promoters (red dots) they increase at low SA origin-distal promoters (blue dots).

**Figure 4.**
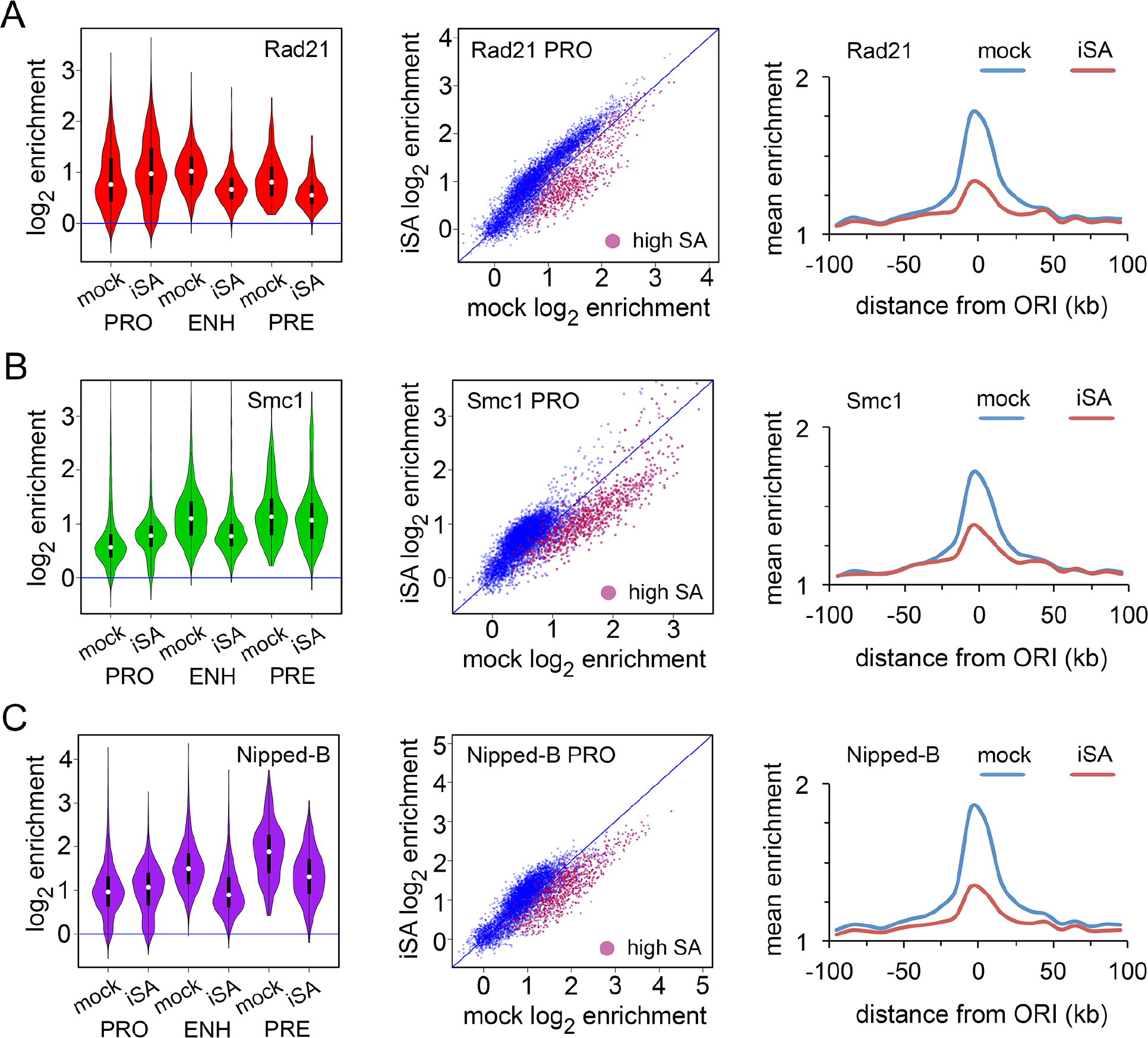
SA targets Rad21, Smc1 and Nipped-B to enhancers and origin-proximal promoters in BG3 cells. An example of SA protein depletion is in Supplemental Fig. S3A. (*A*) Effects of SA depletion (iSA) on Rad21 localization. Violin plots (left) show the distribution of Rad21 enrichment at promoters (PRO) enhancers (ENH) and PREs (PRE) in mock control cells and cells depleted for SA. Promoter dot plots (middle) show enrichment in mock control cells vs. SA-depleted cells. High SA promoters are plotted in red. Origin meta-analysis (right) shows Rad21 distribution surrounding early replication origins in mock control cells (blue) and SA-depleted (iSA) cells (red). (*B*) Effects of SA depletion on Smc1 location. (*C*) Effects of SA depletion on Nipped-B localization. Median values for all occupancy distributions and Wilcoxon p values for mock vs. SA depletion are in Supplemental Table S2. All occupancy changes are statistically significant except for Smc1 at PREs.

SA depletion reduces total Rad21 protein levels (Supplemental Fig. S3A) and part of the Rad21 decrease at enhancers and origin-proximal promoters may reflect reduced protein levels. Against this idea, Rad21 increases at the origin-distal promoters to levels that are higher than those left on the enhancers, indicating that the remaining amount of total Rad21 does not limit chromosome association. Also, total Nipped-B protein levels are not reduced by SA depletion and Nipped-B also shows a decrease at origin-proximal features and increase at origin-distal promoters. FRAP experiments show that less than 20% of cohesin in the nucleus is bound to chromosomes in *Drosophila* cells indicating that moderate depletion of total cohesin levels does not limit chromosome association (Gause et al. 2010).

Importantly, the opposite effects of SA depletion on Rad21, Smc1 and Nipped-B association with origin-proximal features and origin-distal promoters indicates that epitope masking or poor crosslinking are not responsible for the inability to detect SA at origin-distal promoters. Rad21, Smc1 and Nipped-B all decrease at origin-proximal promoters and enhancers where SA is detected but increase at origin-distal promoters where SA is not detected. If SA were present at origin-distal promoters we would expect Rad21, Smc1 and Nipped-B levels to also decrease upon SA depletion. The Rad21, Smc1 and Nipped-B increases at origin-distal promoters following SA depletion also provide further evidence that cohesin can bind promoters independently of SA.

SA is not part of the cohesin ring (Fig. 1A) and is not required for cohesin to topologically bind chromosomes (Kulemzina et al. 2012). We depleted Smc1 to compare how a ring component influences Nipped-B and cohesin subunit chromosome occupancy (Fig. 5). The Smc1 antibody gives weak signals in western blots so ChIP-seq was used to confirm Smc1 depletion, showing that Smc1 occupancy is globally reduced (Supplemental Table S2, Supplemental Fig. S3B). Supplemental Fig. S4 shows examples of Smc1 reduction at the origin-proximal region containing the *string* gene and an origin-distal region containing *woc*. Smc1 depletion reduces total Rad21 protein to a similar extent as SA depletion (Supplemental Fig. S3A). Smc1 depletion also reduces total Rad21 in human cells (Laugsch et al. 2013).

**Figure 5.**
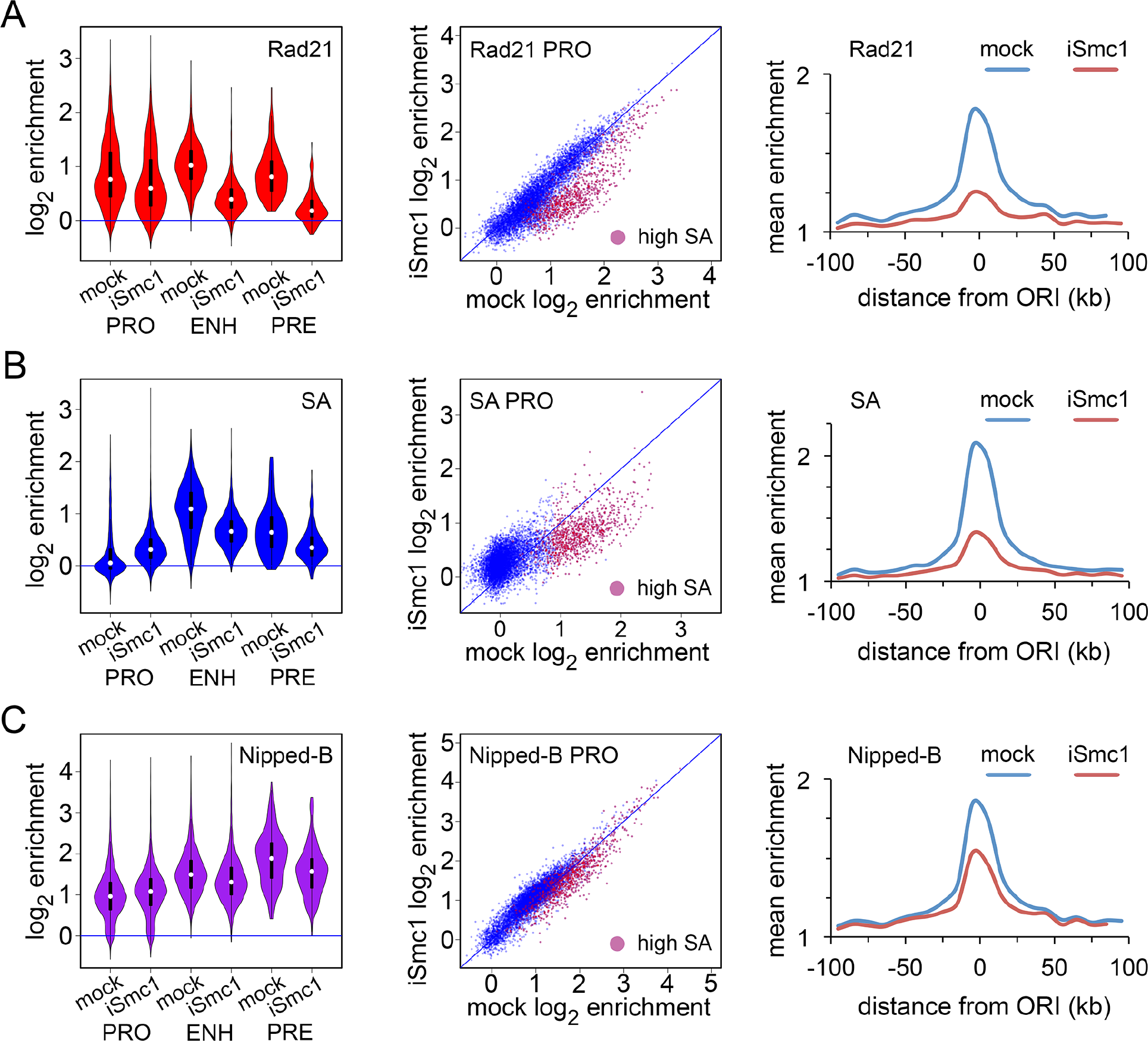
Smc1 facilitates Rad21 and SA association with enhancers and origin-proximal promoters in BG3 cells. Effects of Smc1 depletion (iSmc1) on Smc1 occupancy are shown in Supplemental Fig. S3B. (*A*) Effects of Smc1 depletion on Rad21 localization. (*B*) Effects of Smc1 depletion on SA localization. (*C*) Effects of Smc1 depletion on Nipped-B localization. All occupancy changes are statistically significant (Supplemental Table S2).

Smc1 depletion reduces Rad21 at enhancers, PREs, and origin-proximal promoters (red dots in promoter dot-plots) (Fig. 5A, Supplemental Table S2, Supplemental Fig. S4A). However, as illustrated by promoter dot plot in Fig. 5A (blue dots) and ChIP-seq tracks in Supplemental Fig. S4B, Smc1 depletion has little effect on Rad21 at origin-distal promoters, and the overall Rad21 level at promoters is now higher than at enhancers (violin plot). Some of the Rad21 decrease at the origin-proximal features might reflect reduced total Rad21 protein, but the minimal effect at origin-distal promoters indicates that a moderate reduction in total Rad21 level does not limit chromosome association. The finding that Smc1 depletion reduces Rad21 levels at origin-proximal promoters, where Smc1 is detected, but has little effect at origin-distal promoters indicates that epitope masking or poor crosslinking are not responsible for the sub-proportional Smc1 at origin-distal promoters seen in control cells. These findings also confirm that Rad21 can occupy promoters independently of Smc1 and SA.

SA association with enhancers, PREs, and origin-proximal regions is also Smc1-dependent (Fig. 5B, Supplemental Table S2, Supplemental Fig. S4A). Smc1 depletion decreases SA at high SA promoters and increases SA to modest levels at promoters that normally have no SA (Fig. 5B, Supplemental Fig. S4B). Detection of SA at origin-distal promoters with Smc1 depletion indicates that epitope masking or poor crosslinking does not cause the lack of SA at these promoters in control cells.

Nipped-B associates with promoters independently of Smc1. Smc1 depletion causes statistically significant reductions in Nipped-B at enhancers, PREs, and origin-proximal promoters with slight increases at origin-distal promoters (Fig. 5C, Supplemental Table S2, Supplemental Fig. S4).

Supplemental Fig. S5 shows that Rad21 depletion reduces Nipped-B association with origin-proximal features including enhancers and high SA promoters, and increases Nipped-B at most promoters. All changes are statistically significant (Supplemental Table S2). We conclude that Nipped-B enhancer occupancy depends on SA and Rad21 while association with most promoters is cohesin-independent.

### The MED30 Mediator subunit and the Fs(1)h BET domain protein co-localize with Nipped-B and cohesin

The above experiments show that SA promotes Nipped-B and cohesin occupancy of features close to early origins and that Nipped-B and Rad21 bind origin-distal promoters independently of SA and Smc1. This implies that enhancer and promoter factors differentially recruit Nipped-B, SA and cohesin. The Mediator complex that regulates transcription (Allen and Taatjes 2015) and the BRD4 BET domain protein that binds acetylated histones (Hsu and Blobel 2017) are candidates for such factors. Mammalian Mediator interacts with NIPBL (Nipped-B) (Kagey et al. 2010) and an affinity chromatography-mass spectrometry screen revealed that the *Drosophila* MED30 Mediator subunit interacts with Nipped-B (Guruharsha et al. 2012). It was recently reported that human BRD4 interacts with NIPBL and that *BRD4* mutations cause birth defects similar to those caused by *NIPBL* mutations (Olley et al. 2018).

We performed ChIP-seq for MED1, MED30 and Fs(1)h, the *Drosophila* ortholog of BRD4, to compare them to cohesin. Fig. 6A shows the origin-proximal *string* locus and Fig. 6B shows an origin-distal region. MED30 and Fs(1)h spread similarly to Nipped-B at the *string* enhancers and MED1 displays more distinct peaks (Fig. 6A). MED30 and Fs(1)h are high at enhancers (Fig. 6C) and strongly origin-centric (Fig 6D). MED1 is less origin-centric, similar to RNA polymerase (Rpb3). The origin-centric distribution of MED30 and Fs(1)h led us to test how they influence Nipped-B association with enhancers and promoters.

**Figure 6.**
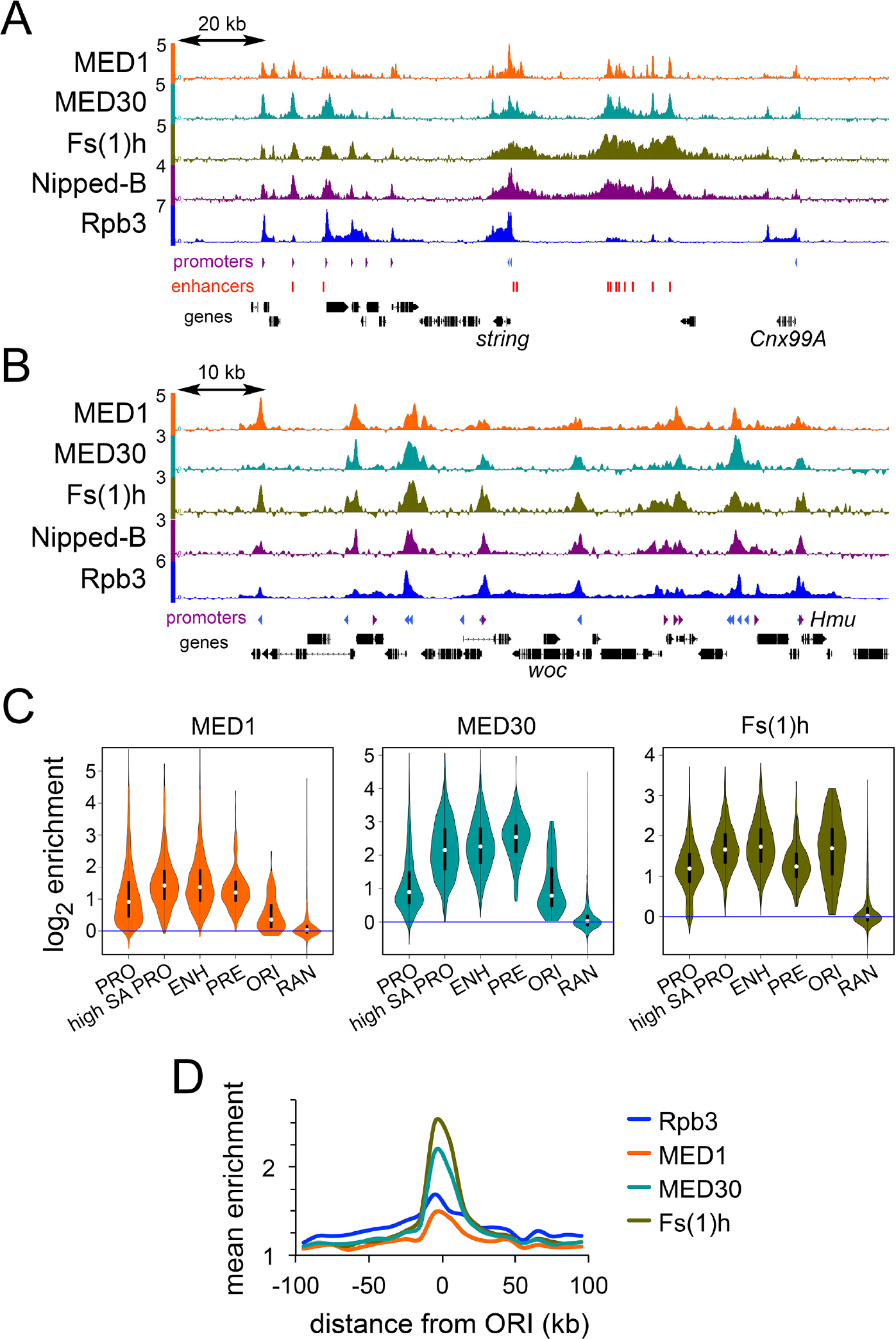
The MED30 Mediator subunit and Fs(1)h BET domain protein co-localize with Nipped-B and are origin-centric in BG3 cells. (*A*) ChIP-seq for the indicated proteins in an origin-proximal region containing the *string* (*cdc25*) gene. (*B*) ChIP-seq for the indicated proteins in an origin-distal region containing the *woc* gene. (*C*) Violin plot distributions for MED1 (orange) MED30 (cyan) and Fs(1)h (olive) occupancy at all active promoters (PRO) SA-occupied promoters (high SA PRO) enhancers (ENH) PREs (PRE) centers of early DNA replication origins (ORI) and randomly positioned sequences (RAN). (*D*) Meta-origin analysis of Rpb3 (blue) MED1 (orange) MED30 (cyan) and Fs(1)h (olive) occupancy.

### MED30 facilitates Nipped-B association with promoters and Fs(1)h promotes Nipped-B enhancer occupancy

MED30 depletion decreases Nipped-B at all promoters with slight increases at enhancers and PREs (Fig. 7A, Supplemental Table S2). Thus, although MED30 is at both promoters and enhancers, it facilitates Nipped-B association only at promoters, suggesting that other factors influence the ability of MED30 to recruit Nipped-B.

**Figure 7.**
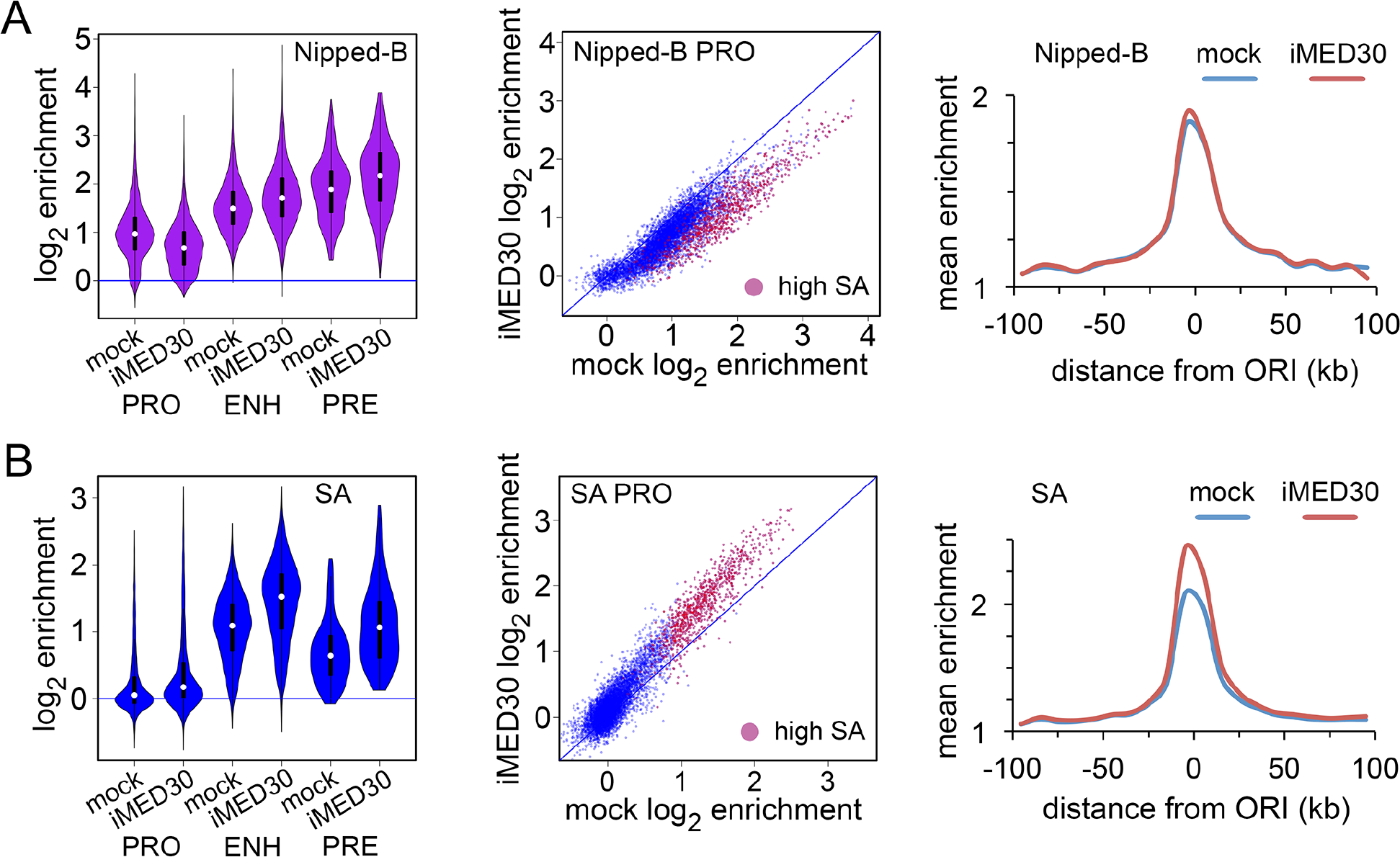
The MED30 Mediator subunit facilitates Nipped-B association with promoters in BG3 cells. An example of MED30 protein depletion is in Supplemental Fig. S3A. (*A*) Effects of MED30 depletion (iMED30) on Nipped-B localization. (*B*) Effects of MED30 depletion on SA localization. All changes in Nipped-B and SA occupancy are statistically significant except at the centers of early origins (Supplemental Table S2).

MED30 depletion increases SA at all features (Fig. 7B, Supplemental Table S2). and SA depletion globally increases MED30 occupancy (Supplemental Fig. S6A, Supplemental Table S2) suggesting that SA and MED30 compete for binding. Smc1 depletion slightly reduces MED30 at all features (Supplemental Fig. S6B) indicating that the SA – MED30 competition is specific.

Fs(1)h promotes Nipped-B association with enhancers and origin-proximal promoters (Fig. 8A, Supplemental Table S2). We treated cells with the JQ1 inhibitor of BET domain binding to acetylated histones (Filippakopoulos et al. 2010) for eight hours to globally reduce Fs(1)h binding (Supplemental Fig. S7A). JQ1 treatment reduces Nipped-B at enhancers, high SA promoters and PREs with an overall decrease in origin-proximal regions. There is little effect at origin-distal promoters (Fig. 8A).

**Figure 8.**
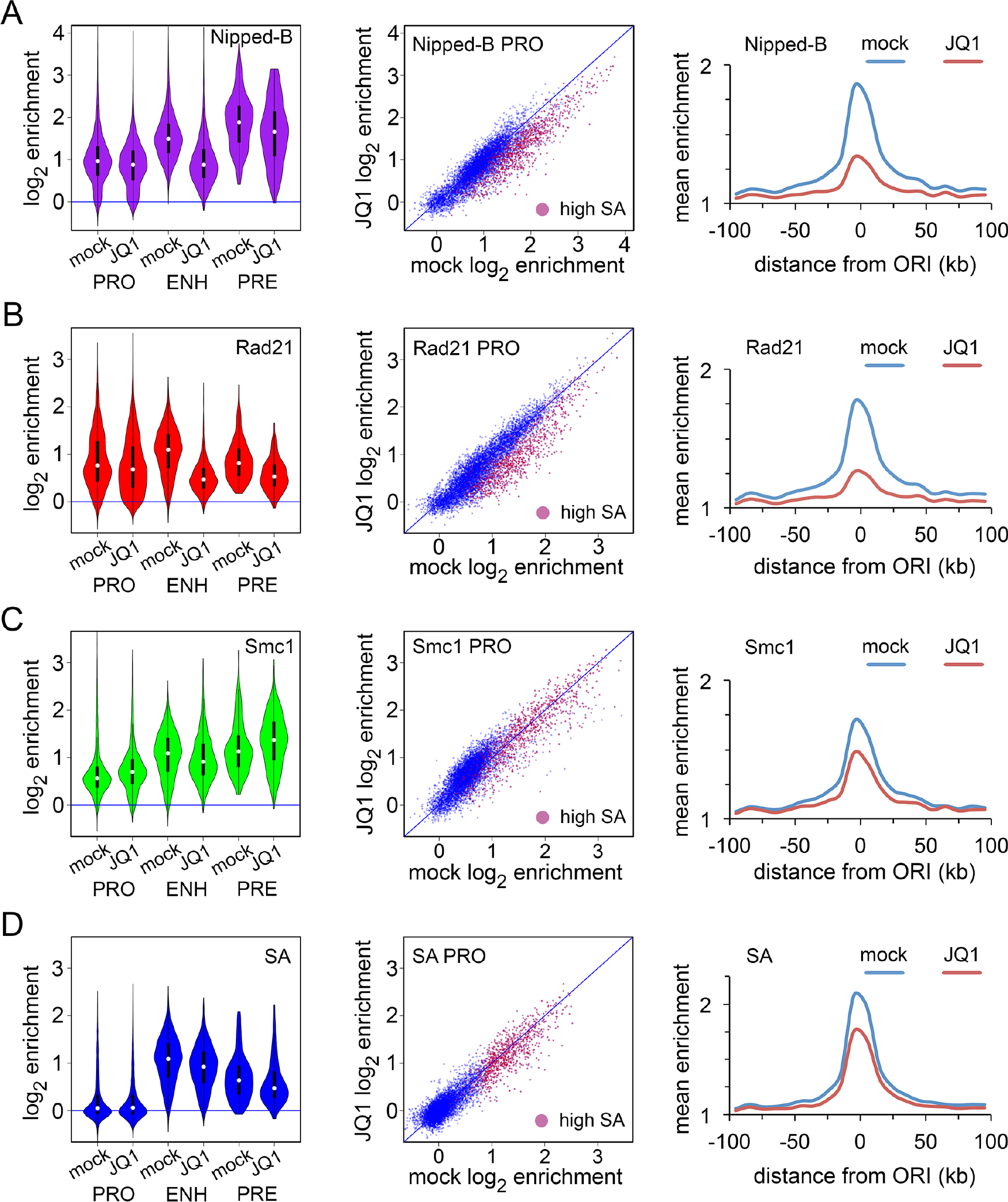
The Fs(1)h BET domain protein promotes association of Nipped-B and Rad21 with enhancers and origin-proximal promoters. The effects of the JQ1 inhibitor on Fs(1)h binding and cell cycle are shown in Supplemental Fig. S6. (*A*) Effects of JQ1 on Nipped-B occupancy. (*B*) Effects of JQ1 on Rad21 occupancy. (*C*) Effects of JQ1 on Smc1 occupancy. (*D*) Effects of JQ1 on SA occupancy. Effects of JQ1 on Nipped-B occupancy are statistically significant except at PREs (Supplemental Table S2). All effects on Rad21 occupancy are significant. Effects on Smc1 occupancy are significant except at high SA promoters. Changes in SA occupancy are significant except at promoters and PREs.

The effects of JQ1 treatment on Rad21 are similar to the effects on Nipped-B, with decreases at enhancers, high SA origin-proximal promoters, and PREs and little effect at origin-distal promoters (Fig. 8B, Supplemental Table S2). Smc1 slightly increases at origin-distal promoters and many PREs, and decreases at enhancers with little change at high SA promoters (Figure 8C, Supplemental Table S2). JQ1 slightly reduces SA at high SA promoters, enhancers and PREs, with little effect at most promoters (Figure 8D, Supplemental Table S2). JQ1 modestly reduces MED30 at enhancers, with even smaller effects at other features (Supplemental Table S2, Supplemental Fig. S6C).

The picture that emerges is that Fs(1)h facilitates Nipped-B and Rad21 association with enhancers, but only slightly influences SA and Smc1 occupancy. This suggests that SA and Smc1 can be recruited independently of Nipped-B and Rad21 to enhancers. JQ1 stops most cells in the G2 phase of the cell cycle (Supplemental Fig. S7B). Thus, some effects on cohesin distribution could reflect differences between populations in which roughly half the cells are in G2 (control) as opposed to the majority (JQ1). This might explain minor changes in SA and Smc1, but seems unlikely to cause the dramatic Nipped-B and Rad21 decreases at enhancers. Indeed, as described below, *Nipped-B*, *Rad21*, and *fs(1)h* mutations show genetic interactions during development when cell division is not blocked.

### *Nipped*-B and *Rad21* (*vtd*) mutations interact genetically with the *fs(1)h*^*1*^ hypomorphic mutation

We tested the in vivo significance of Fs(1)h effects on Nipped-B and Rad21 localization using the hypomorphic *fs(1)h*^*1*^ mutation. *fs(1)h*^*1*^ was recovered in a screen for female-sterile mutations on the X Chromosome (Gans et al. 1975). Null *fs(1)h* alleles are lethal but *fs(1)h*^*1*^ is a viable missense mutation (Digan et al. 1986; Florence and Faller 2008).

*Nipped-B* and *Rad21* (*vtd*) mutations dominantly reduce *fs(1)h*^*1*^ viability (Fig. 9A). At 29°, 62% of the expected *fs(1)h^1^/Y* males were recovered relative to their *fs(1)h^1^/+* sisters. The heterozygous *Nipped-B*^*407*^ null mutation does not reduce viability in wild-type backgrounds (Rollins et al. 1999; Wu et al. 2015) but reduces viability of *fs(1)h^1^/Y* males to 5% of their *fs(1)h^1^/+; Nipped-B^407^/+* sisters. *Nipped-B^NC41^*, a truncation mutation (Gause et al. 2008) dominantly reduces *fs(1)h*^*1*^ male viability to 25%. *vtd^4^*, an partial *Rad21* deletion (Hallson et al. 2008) dominantly reduces *fs(1)h*^*1*^ male viability to 12%, and *vtd^ex15^*, another partial deletion (Pauli et al. 2008) reduces viability to 49%. All viability reductions are statistically significant, except with *vtd^ex15^*.

**Figure 9.**
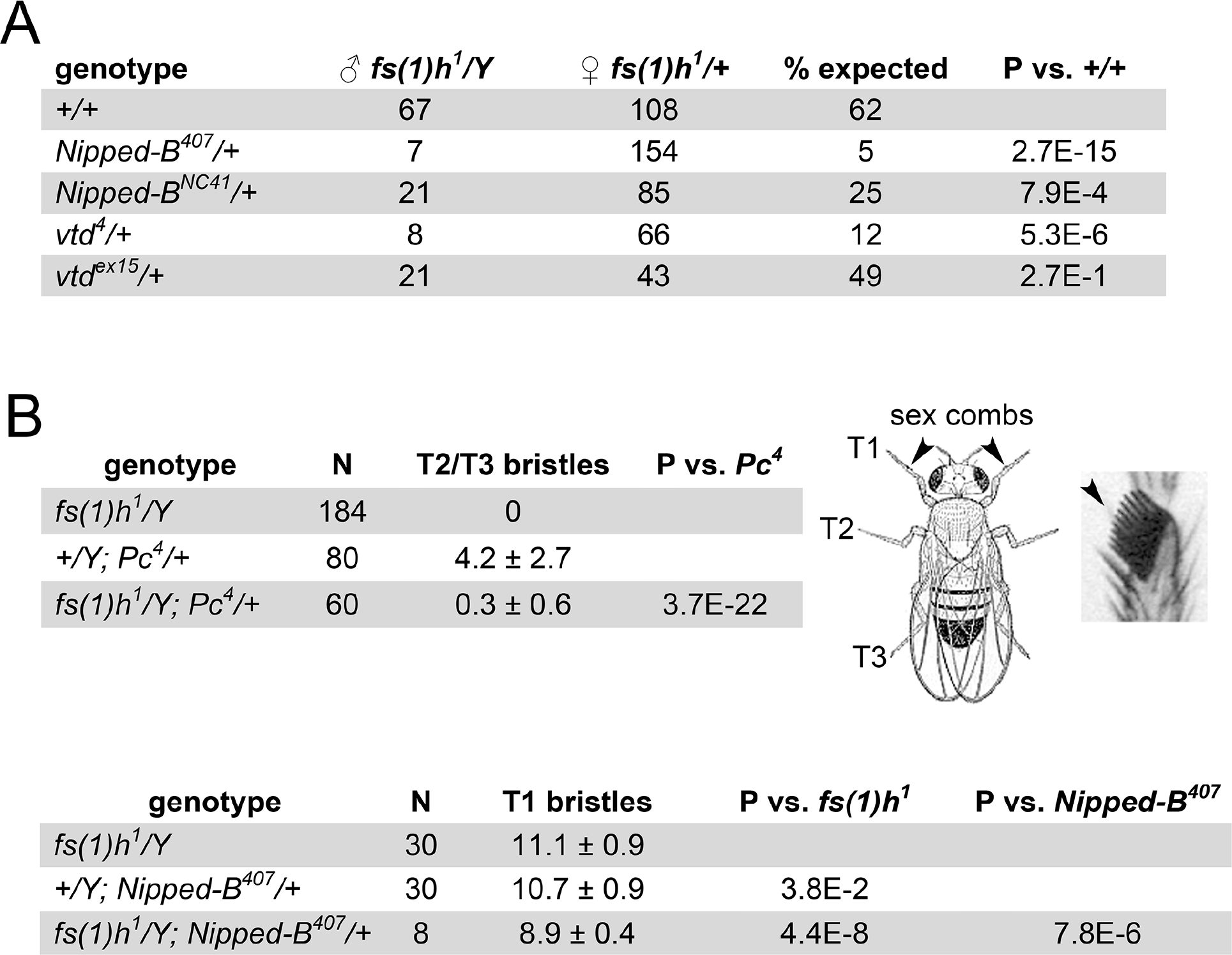
*Nipped-B* and *Rad21* (*vtd*) mutations dominantly enhance *fs(1)h*^*1*^ mutant phenotypes. Crosses were conducted at 29°. (*A*) *Nipped-B* and *Rad21* (*vtd*) mutations dominantly decrease *fs(1)h*^*1*^ male viability. The numbers of *fs(1)h^1^/Y* males with the indicated genotypes and their *fs(1)h^1^/+* sisters recovered are given. The % expected is the male to female ratio. P values were calculated using Fisher’s exact test. (*B*) *fs(1)h*^*1*^ suppresses the ectopic T2 and T3 leg sex comb bristles in *Pc*^*4*^ mutant males and reduces the number of T1 sex comb bristles when combined with heterozygous *Nipped-B^407^*. The diagram shows a male fly indicating the T1, T2, and T3 legs and a magnified view of T1 sex comb bristles. The tables give the number of legs scored (N) with the average number of bristles per leg and the standard deviation. P values were calculated using the t-test.

*Nipped-B* and *Rad21* (*vtd*) mutations dominantly suppress homeotic transformations caused by *Pc* mutations (Kennison and Tamkun 1988; Hallson et al. 2008; Schaaf et al. 2013b). *fs(1)h*^*1*^ similarly suppresses ectopic sex comb bristles on the T2 and T3 legs of *Pc^4^/+* males (Fig. 9B). Strikingly, combining *fs(1)h*^*1*^ with heterozygous *Nipped-B*^407^ reduces the number of T1 sex comb bristles in the surviving males (Fig. 9B). The genetic interactions between *fs(1)h*, *Nipped-B* and *Rad21* indicate that the proteins they encode function together in vivo.

## Discussion

Our experiments show that SA helps recruit cohesin complexes to enhancers, which are all located close to early DNA replication origins (Fig. 10A) and to those promoters that are also close to origins. In contrast to SA, Nipped-B and Rad21 also occupy origin-distal promoters (Fig. 10A). We posit that at these promoters, Smc1-Smc3-Rad21 tripartite rings sometimes exchange for the Rad21 in Nipped-B–Rad21 complexes with subsequent loading of cohesin lacking SA.

**Figure 10.**
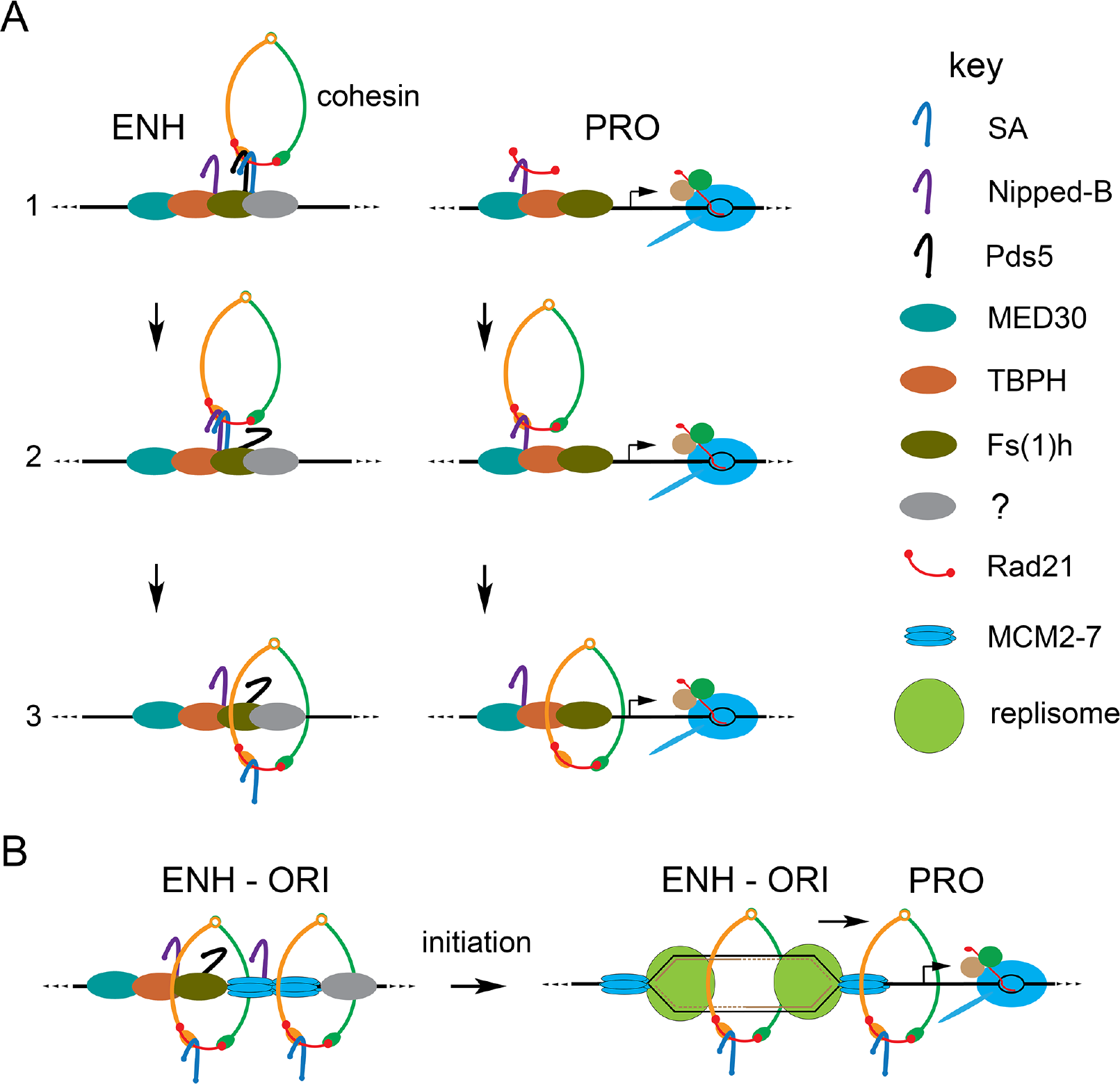
Theoretical models for cohesin recruitment and the roles of origins and DNA replication in sister chromatid cohesion and enhancer-promoter communication. (*A*) Cohesin recruitment to enhancers (ENH) and promoters (PRO). The factor key is on the right. At enhancers (left) we posit that that Pds5 (black) and SA (blue) recruit tripartite cohesin rings (step 1) and that Nipped-B (purple) displaces Pds5 (step 2) to load cohesin topologically (step 3). SA association with enhancers is facilitated by Pds5 (Misulovin et al 2018) and unknown enhancer-specific proteins (gray). At promoters (right) we envision that the MED30 Mediator subunit (cyan) and the TBPH protein (orange) (Swain et al. 2016) recruit Nipped-B and Rad21 (red) without Smc1-Smc3 dimers or SA (step 1). At some frequency, Nipped-B – Rad21 complexes capture Smc1-Smc3 dimers or exchange Rad21 for Smc1-Smc3-Rad21 rings (step 2) and SA-deficient cohesin is loaded (step 3). (*B*) We theorize that enhancers capture translocating MCM2-7 helicase complexes (Powell et al. 2015) to position early replication origins (left). DNA unwinding by MCM2-7 upon initiation of replication topologically captures both single-stranded templates within cohesin rings behind the nascent forks to establish sister chromatid cohesion (right). Replication forks push other SA-containing cohesin rings to be captured by neighboring promoters, facilitating enhancer-promoter communication (right).

In contrast to the relatively low levels of SA-deficient cohesin at origin-distal promoters, cohesin contains SA at enhancers and origin-distal promoters and is present at higher levels. The evidence that cohesin lacks SA at most origin-distal promoters, and that SA also binds chromosomes independently of cohesin, leads us to now consider SA to be a cohesin regulator that controls cohesin localization instead of a cohesin subunit.

Our experiments also show that in addition to SA, the Fs(1)h (BRD4) mitotic bookmarking protein facilitates cohesin association with enhancers and the origin-proximal promoters (Fig. 10A). Genetic evidence confirms that Fs(1)h functions together with Nipped-B and Rad21 in vivo to support development. In contrast to the origin-centric roles for SA and Fs(1)h, the MED30 subunit of the Mediator complex facilitates association of Nipped-B and Rad21 with active promoters, but not with enhancers.

### Enhancer-promoter communication

Only promoters that are close to enhancers and origins are occupied by high levels of cohesin containing SA. We theorize that these are the promoters that are targeted by enhancers. We envision that DNA replication pushes cohesin from enhancers to origin-proximal promoters (Fig. 10B) based on the evidence that replication origins form preferentially at enhancers, and prior indications that replication pushes cohesin (Kanke et al. 2016; Misulovin et al. 2018). We do not know if the Nipped-B and Rad21 that bind origin-distal promoters independently of SA and Smc1 (Fig. 10A) influence gene transcription. This will be challenging to unravel because Nipped-B and Rad21 are essential for complete cohesin rings to bind to chromosomes.

Since it was discovered that sister chromatid cohesion proteins facilitate expression of enhancer-activated genes (Rollins et al. 1999) it has been proposed that enhancer-promoter looping could be supported by intra-chromosomal cohesion. In the simplest version, a cohesin ring topologically encircles DNA near both the enhancer and the promoter to hold them together. The cohesin at the enhancer and promoter are thus the same molecules. Some of our findings argue against this idea. In particular, MED30 depletion reduces Nipped-B and Rad21 at origin-proximal promoters but not at the linked enhancers, indicating that different cohesin molecules are present at the enhancers and promoters. It could be that a cohesin ring at a promoter interacts with another at an enhancer to handcuff them together, or that cohesin interacts with Mediator, BRD4 or other proteins to stabilize enhancer-promoter looping.

Cohesin is removed from chromosomes at mitosis and loaded in early G1. Thus, the idea that DNA replication localizes cohesin to facilitate enhancer-promoter communication raises the question of how cohesin supports enhancer function in G1 before replication. One idea is that mitotic bookmarking factors facilitate cohesin loading at enhancers and target promoters. The BRD4 ortholog of Fs(1)h remains bound to mitotic chromosomes and promotes rapid reactivation of transcription after cell division (Dey et al. 2000, Zhao et al. 2011). Thus, the finding that inhibiting Fs(1)h chromosome binding reduces Nipped-B and Rad21 at enhancers and origin-proximal promoters without going through cell division supports the idea that Fs(1)h marks them for cohesin loading.

### Sister chromatid cohesion

We hypothesize that origins form at enhancers because enhancers trap the sliding MCM2-7 helicase that will initiate DNA replication (Fig. 10B). Localization of cohesin to enhancers and origins suggests a simple model for how sister chromatid cohesion is established. Upon initial unwinding of the DNA template by MCM2-7, cohesin behind the nascent replication forks encircles the two single-stranded templates, passively establishing cohesion while cohesin in front of the forks is pushed to origin-proximal promoters (Fig. 10B).

This model explains why Pds5, a cohesin removal factor, and SA, which is not required for cohesin to bind chromosomes topologically (Kulemzina et al. 2012) are required for sister chromatid cohesion. By positioning cohesin at enhancers they ensure that the nascent sister chromatids will be topologically trapped within cohesin (Fig. 10B). This does not require that replisomes move through cohesin or new cohesin loading behind the fork as proposed in other models (Uhlmann 2016, Villa‐Hernández and Bermejo 2018). It is consistent with the finding that cohesin can remain chromosome-bound and establish cohesion during DNA replication in the absence of the Wapl removal factor (Rhodes et al. 2017).

### Parallels with vertebrate cohesin

Mammals have two SA orthologs, SA1 (STAG1) and SA2 (STAG2). Only SA2-containing cohesin is present at enhancers in human cells (Kojic et al. 2018) suggesting that SA2 is the functional ortholog of *Drosophila* SA. SA2 binds DNA independently of cohesin in vitro with a preference for single-stranded DNA and structures resembling replication forks (Countryman et al. 2018). This is consistent with our findings that SA is origin-centric and spreads further than cohesin around enhancers.

Mutations in the *STAG2* gene encoding SA2 cause intellectual and growth deficits overlapping those seen in cohesinopathies caused by mutations in *NIPBL* or cohesin subunit genes (Mullegama et al. 2017, Soardi et al. 2017, Mullegama et al. 2018, Yuan et al. 2018). Individuals with *BRD4* mutations display similar birth defects, and BRD4 and NIPBL co-localize at enhancers (Olley et al. 2018). These studies agree with our findings that SA and Fs(1)h facilitate association of Nipped-B and Rad21 with enhancers and that Fs(1)h and Nipped-B function together in development.

Our data show parallels with cohesin loading in *Xenopus*. Cohesin loading in *Xenopus* oocyte extracts requires assembly of the pre-replication complex that licenses replication origins, and the Cdc7-Drf1 kinase that activates the pre-replication complex interacts with NIPBL (Gillespie and Hirano 2004; Takahashi et al. 2004; Takahashi et al. 2008). This places cohesin at the site of replication initiation, similar to the role of SA in *Drosophila*.

Specialized DNA replication factors are needed to establish sister chromatid cohesion in yeast (Skibbens 2009) but it is unclear whether they are required at progressing forks or only upon initiation of replication. A study in human cells showed that NIPBL and cohesin interact with the MCM2-7 helicase (Zheng et al. 2018). The authors suggest that NIPBL bound to MCM2-7 is transiently held by the replisome and transferred behind the fork to load cohesin and establish sister cohesion, but it is possible that interactions with NIPBL could also trap MCM2-7 at enhancers prior to replication. Whether or not recruiting both MCM2-7 and cohesin to origins is sufficient to establish cohesion or whether cohesion requires new cohesin loading behind the replication fork remains to be resolved.

## Materials and methods

### Cell culture, RNAi depletion and JQ1 treatment

ML-DmBG3-c2 (BG3) cells were cultured and proteins were depleted by RNAi as described (Schaaf et al. 2009). Cells were treated with 10 μM JQ1 in the medium for 8 hours to inhibit Fs(1)h binding and cell cycle stages were determined by propidium iodide staining and FACS analysis in the Saint Louis University Flow Core.

### ChIP-seq and quantification

ChIP-seq was performed and quantified as detailed elsewhere (Dorsett and Misulovin 2017) using concurrent experiments, overlapping sets of chromatin preparations, multiple biological repeats and validated antibodies (Supplemental Methods, Supplemental Table S1).

Enhancers, PREs and active promoters were identified and defined as 500 bp sequences based on DNaseI hypersensitivity, histone modifications (H3K4me1, H3K4me3, H3K27me3, H3K27ac) and PRO-seq data (Schaaf et al. 2013a, Schaaf et al. 2013b, Swain et al. 2016, Pherson et al. 2017, Misulovin et al. 2018). Occupancy of individual features was calculated using bed files and scripts provided in prior publications (Swain et al. 2016; Pherson et al. 2017; Misulovin et al. 2018). Details are in Supplemental Methods.

### Early S phase DNA replication

Early S phase DNA replication was quantified by adapting an origin-mapping protocol (MacAlpine et al. 2004) with EdU thymidine analog detection of newly-synthesized DNA (Ramachandran and Henikoff 2016). Details are in Supplemental Methods.

### MED1 and MED30 antibodies

A His(6) fusion to the 1140-1475 C terminal residues of MED1 was expressed in *E. coli*, purified by nickel chromatography under denaturing conditions, and insoluble protein was used to immunize a rabbit at Josman, LLC (Napa, CA). A His(6) fusion to full length MED30 (residues 1-318) was expressed in *E. coli*, purified by nickel chromatography, and the insoluble protein used to immunize a guinea pig at Pocono Rabbit Farm and Laboratory (Canadensis, PA). Antibody specificities were confirmed by western blots of whole cell extracts of control and RNAi-depleted BG3 cells (Supplemental Fig. S3A).

### Genetic crosses

*Drosophila* stocks were maintained and crosses conducted as described (Rollins et al. 1999). *fs(1)h^1^* stocks were obtained from the Bloomington *Drosophila* Stock Center at Indiana University. *Nipped-B*, cohesin and *Pc* mutant stocks are described previously (Rollins et al. 1999; Gause et al. 2008; Schaaf et al. 2013b). Fisher’s exact test and t-tests were used as indicated in Fig. 9.

### Data access

ChIP-seq data are available from GEO: GSE118484.

## Acknowledgements

We thank Igor Dawid (NICHD) for Fs(1)h antibodies, Fran Sverdrup (Saint Louis University) for JQ1 and Srinivas Ramachandran (University of Colorado) for helpful discussions about the EdU experiments. We thank Frank Uhlmann (Francis Crick Institute) and Hongtao Yu (HHMI, UT Southwestern) for helpful comments on the manuscript. This work was supported by grants from the NIH (R01 GM108714) to DD and the CdLS Foundation to MP.

## Disclosure declaration

The authors have no conflicts of interest to disclose.

