## Supplemental Material for "Cohesin occupancy and composition at enhancers and promoters are linked to DNA replication origin proximity in *Drosophila*"

**Supplemental Methods** (pages 2-3)

**Supplemental Table S1.** ChIP-seq replicates and genome-wide Pearson correlations. (page 4)

**Supplemental Table S2.** Median values of Rad21, SA, Smc1, Nipped-B, MED30, and Fs(1)h enrichment at promoters, enhancers, PREs and early replication origin centers in control and treated BG3 cells with statistical tests. (page 5)

**Supplemental Figure S1.** ChIP-seq with preimmune serum in BG3 cells. (page 6)

**Supplemental Figure S2.** Early DNA replication origins align with clusters of enhancers and SA depletion selectively reduces Rad21, Smc1 and Nipped-B association with enhancers and origin-proximal promoters in BG3 cells. (page 7)

**Supplemental Figure S3.** Effects of RNAi depletions on protein levels or chromosome association. (page 8)

**Supplemental Figure S4.** Smc1 depletion selectively reduces cohesin levels on enhancers and origin-proximal promoters in BG3 cells. (page 9)

**Supplemental Figure S5.** Rad21 facilitates Nipped-B and SA association with enhancers, high SA promoters and origin-proximal regions in BG3 cells. (page 10)

**Supplemental Figure S6.** SA inhibits MED30 association with functional features in BG3 cells. (page 11)

**Supplemental Figure S7**. Effects of JQ1 on Fs(1)h chromosome association and cell cycle in BG3 cells. (page 12).

### Supplemental methods

#### ChIP-seq and quantification

ChIP-seq was performed as described (Dorsett and Misulovin 2017) using SA, Rad21, Smc1 and Nipped-B antibodies previously validated for specificity and use in ChIP-seq (Dorsett et al. 2005; Gause et al. 2008; Misulovin et al. 2008; Gause et al. 2010; Schaaf et al. 2013a; Schaaf et al. 2013b; Swain et al. 2016; Misulovin et al. 2018). All experiments used the same antibody bleed for each protein. Fs(1)h antibodies recognizing both the short and long forms were obtained from Igor Dawid (NICHD) and validated by ChIP-seq after JQ1 treatment (Supplemental Fig. S6). MED1 and MED30 antibodies were made and validated as described in the Materials and Methods.

ChIP-seq base pair coverage was normalized to input chromatin base pair coverage using sliding 250 bp windows to generate enrichment values every 50 bp (Dorsett and Misulovin 2017). This method shows no significant enrichment using preimmune serum (Fig. 1, Supplemental Fig. S1). Each ChIP-seq replicate was sequenced to at least 10-fold genome coverage and normalized to input chromatin sequenced to greater than 50-fold genome coverage.

ChIP-seq replicates were tested for their genome-wide correlation and averaged before calculating occupancy of genomic features. There was a minimum genome-wide Pearson correlation of 0.6 across all 2.56 million data points for all replicates (Supplemental Table S1). Occupancy of individual features was calculated as the arithmetic mean of the individual enrichment data points spaced 50 bp apart within each 500 bp feature. Bed files for each feature type and scripts for calculating occupancy are provided in prior publications (Swain et al. 2016; Pherson et al. 2017; Misulovin et al. 2018). Features in heterochromatin were removed to reduce distortions caused by repetitive sequences. The threshold tool in the Integrated Genome Browser (Freese et al. 2016) was used to identify regions of SA enrichment in the 95$^{th}$ percentile over regions ≥300 bp and BEDTools (Quinlan and Hall 2010) was used to identify promoters that overlap these regions. P values comparing controls to experimental samples were calculating using the Wilcoxon test of medians (Supplemental Table S2). For the meta-analyses, enrichment in each bin was calculated as a harmonic mean instead of an arithmetic average to minimize distortion by outliers with particularly high enrichment. Overlapping features were excluded to reduce distortions.

#### Early S phase DNA replication

BG3 cells were cultured in medium containing 1 mM hydroxyurea for 3 hours before adding the EdU (ThermoFisher) thymidine analog at 10 micromolar final concentration in the medium overnight (~16 hours). Cells were removed from the plates by scraping and fixed with 1% formaldehyde (20 mL per 100 million cells) for 12 min at room temperature. Fixation was stopped by addition of glycine to 0.25 M final concentration for 5 min. Cells were collected by centrifugation, washed twice with PBS and stored as a pellet at -80$^{o}$.

To attach biotin to the EdU residues, the cell pellets were permeabilized in 2 mL of PBS containing 0.25% Triton X-100 for 30 min at room temperature, collected by centrifugation, and washed twice with PBS containing 0.5% bovine serum albumin. The washed cell pellet was suspended in 920 microliters of PBS, followed by addition of biotin-azide (Azide-PEG3-Biotin Conjugate, Millipore Sigma, stock concentration 2 mM in DMSO) to a final concentration of 100 micromolar, sodium L-ascorbate (fresh 1 M stock concentration) to 10 mM final concentration, and $CuSO_4$ (100 mM stock concentration) to 2 mM final concentration in a total volume of 1 mL before incubation in the dark at room temperature for 60 min. The cells were collected by centrifugation and washed three times with 6 mL of PBS.

Chromatin was prepared from the washed cells as described (Dorsett and Misulovin 2017) except the cells were suspended in 1 mL sonication buffer per 100 million cells, the sonication buffer contained 0.4% (w/v) sodium dodecyl sulfate (SDS) and sonication was conducted for 5 cycles. The sonicated suspension was adjusted to 0.5% lauryl sarcosine, mixed by rotation for 10 min at room temperature and clarified by centrifugation at 15,000 g for 10 min.

DNA was isolated from the chromatin after reversing the crosslinks. One mL of chromatin was diluted with 1 mL of TE (10 mM Tris.HCl pH 7.9, 1 mM EDTA) and 40 microliters of RNAse A (4 milligrams per mL) was added before incubation at 37$^o$ for 30 min. Proteinase K (50 microliters of 20 milligrams per mL) and 50 microliters of 20% SDS were added before incubation at 65$^o$ overnight. After the incubation, phenol:chloroform:isoamyl alcohol (25:24:1, pH 7.5) extraction was performed using phase lock gel tubes (QuantaBio 5 Prime Phase Lock Gel Light) and the DNA was purified using columns (Qiagen QIAquick PCR Purification). DNA was eluted from the columns twice with 55 microliters of the provided elution buffer.

EdU-containing DNA was isolated from the purified DNA using magnetic streptavidin beads (Dynabeads® M-280 Streptavidin, ThermoFisher). Twenty microliters of beads were washed twice before use with 1 mL of WB (5 mM Tris.HCl pH 7.5, 0.5 mM EDTA, 1 M NaCl, 0.1% Tween 20). 100 microliters of DNA was diluted with 100 microliters of 2X WB and incubated with the streptavidin beads for 30 min at room temperature. Beads were collected with a magnet and washed 3 times with 1 mL WB for 5 min. DNA was eluted from the beads by suspending in 400 microliters of 0.3X TE containing 1% SDS and 0.5 micrograms per mL Proteinase K and incubating at 65$^o$ for 20 min. The bead suspension was extracted with 400 microliters of phenol:chloroform:isoamyl alcohol (25:24:1, pH 7.5) and DNA was purified from the supernatant using a column (Qiagen MinElute PCR Purification). Sequencing libraries were constructed from the input and EdU-purified DNA using the Ion Torrent Fragment Library Kit (ThermoFisher) according to the manufacturer's protocol. After sequencing, enrichment of EdU-containing DNA coverage relative to the input coverage was calculated as per the ChIP-seq protocol (Dorsett and Misulovin 2017) with a sliding window of 2,450 bp. Quantification of enrichment at functional features was performed as described above for the ChIP-seq experiments.

| | | Pearson correlations | | | | | |
|---|---|---|---|---|---|---|---|
| **ChIP-seq replicate** | **chromatin** | **1-2** | **1-3** | **1-4** | **2-3** | **2-4** | **3-4** |
| Fs(1)h JQ1 1 | 145 | | | | | | |
| Fs(1)h mock 1 | 161.1 | | | | | | |
| Fs(1)h mock 2 | 161.2 | 0.87 | | | | | |
| MED1 mock 1 | 60 | | | | | | |
| MED1 mock 2 | 58 | 0.89 | | | | | |
| MED30 JQ1 1 | 145 | | | | | | |
| MED30 JQ1 2 | 146 | 0.80 | | | | | |
| MED30 mock 1 | 161.2 | | | | | | |
| MED30 mock 2 | 164 | 0.82 | | | | | |
| MED30 iSA 1 | 160.1 | | | | | | |
| MED30 iSA 2 | 160.2 | 0.90 | | | | | |
| MED30 iSmc1 1 | 186.1 | | | | | | |
| MED30 iSmc1 2 | 186.2 | 0.86 | | | | | |
| Nipped-B JQ1 1 | 146 | | | | | | |
| Nipped-B JQ1 2 | 171 | 0.61 | | | | | |
| Nipped-B mock 1 | 161.1 | | | | | | |
| Nipped-B mock 2 | 164 | 0.73 | | | | | |
| Nipped-B iMED30 1 | 181 | | | | | | |
| Nipped-B iMED30 2 | 182 | 0.73 | | | | | |
| Nipped-B iMED30 3 | 183 | | 0.74 | | 0.73 | | |
| Nipped-B iRad21 1 | 159.1 | | | | | | |
| Nipped-B iRad21 2 | 159.2 | 0.69 | | | | | |
| Nipped-B iSA 1 | 160.1 | | | | | | |
| Nipped-B iSA 2 | 160.2 | 0.62 | | | | | |
| Nipped-B iSmc1 1 | 186.1 | | | | | | |
| Rad21 JQ1 1 | 146 | | | | | | |
| Rad21 JQ1 2 | 171 | 0.60 | | | | | |
| Rad21 mock 1 | 161 | | | | | | |
| Rad21 mock 2 | 164 | 0.63 | | | | | |
| Rad21 iSA 1 | 160.1 | | | | | | |
| Rad21 iSA 2 | 160.2 | 0.67 | | | | | |
| Rad21 iSA 3 | 179 | | 0.69 | | 0.68 | | |
| Rad21 iSmc1 1 | 186.1 | | | | | | |
| SA JQ1 1 | 145 | | | | | | |
| SA JQ1 2 | 146 | 0.69 | | | | | |
| SA mock 1 | 161.2 | | | | | | |
| SA mock 2 | 164 | 0.75 | | | | | |
| SA mock 3 | 164 | | 0.77 | | 0.71 | | |
| SA mock 4 | 165 | | | 0.73 | | 0.69 | 0.69 |
| SA iMED30 1 | 181 | | | | | | |
| SA iMED30 2 | 182 | 0.79 | | | | | |
| SA iMED30 3 | 183 | | 0.78 | | 0.76 | | |
| SA iRad21 1 | 159.1 | | | | | | |
| SA iRad21 2 | 159.2 | 0.61 | | | | | |
| SA iSmc1 1 | 186 | | | | | | |
| Smc1 JQ1 1 | 146 | | | | | | |
| Smc1 JQ1 2 | 171 | 0.60 | | | | | |
| Smc1 mock 1 | 161.1 | | | | | | |
| Smc1 mock 2 | 161.2 | 0.60 | | | | | |
| Smc1 mock 3 | 164 | | 0.60 | | 0.67 | | |
| Smc1 iSA 1 | 160.2 | | | | | | |
| Smc1 iSA 2 | 179 | 0.64 | | | | | |
| Smc1 iSmc1 1 | 186.1 | | | | | | |
| Smc1 iSmc1 2 | 186.2 | 0.61 | | | | | |

**Supplemental Table S1.** ChIP-seq replicates and genome-wide Pearson correlations.

| Protein | Feature (N) | Median $log_2$ enrichment | | | | | | Wilcoxon P values | | | | |
|---|---|---|---|---|---|---|---|---|---|---|---|---|
| | | Mock | iSA | iSmc1 | iRad21 | iMED30 | JQ1 | iSA | iSmc1 | iRad21 | iMED30 | JQ1 |
| Rad21 | PRO (7,398) | 0.76 | ***0.98*** | 0.60 | | | 0.68 | 4.21E-60 | 1.10E-59 | | | 2.95E-28 |
| | High SA PRO (895) | ***1.41*** | ***0.98*** | ***0.62*** | | | ***0.85*** | 2.40E-48 | 9.70E-120 | | | 2.25E-74 |
| | ENH (2,353) | 1.02 | 0.66 | 0.39 | | | 0.47 | 3.65E-202 | 0.00E+00 | | | 0.00E+00 |
| | PRE (195) | 0.81 | 0.55 | 0.18 | | | 0.53 | 2.64E-11 | 1.08E-44 | | | 1.64E-13 |
| | ORI (78) | 0.92 | 0.36 | 0.38 | | | 0.31 | 1.54E-16 | 2.36E-18 | | | 5.64E-18 |
| SA | PRO | 0.05 | | 0.31 | 0.37 | 0.17 | 0.06 | | 0.00E+00 | 0.00E+00 | 1.03E-95 | **9.56E-01** |
| | High SA PRO | 1.25 | | ***0.79*** | ***0.70*** | ***1.68*** | ***1.17*** | | 5.66E-127 | 1.77E-139 | 2.00E-77 | 1.64E-06 |
| | ENH | 1.09 | | 0.66 | 0.58 | 1.52 | 0.93 | | 7.29E-181 | 1.21E-213 | 6.39E-120 | 5.91E-25 |
| | PRE | 0.64 | | 0.35 | 0.40 | 1.07 | 0.47 | | 1.48E-10 | 2.47E-08 | 2.57E-12 | **1.19E-02** |
| | ORI | ***1.38*** | | 0.51 | 0.52 | 1.51 | 0.93 | | 3.86E-19 | 2.38E-19 | **1.68E-02** | 1.70E-06 |
| Smc1 | PRO | 0.57 | 0.78 | 0.45 | | | 0.70 | 2.33E-190 | 3.58E-160 | | | 8.45E-62 |
| | High SA PRO | ***1.55*** | ***1.10*** | ***0.72*** | | | ***1.55*** | 2.44E-55 | 4.69E-150 | | | 2.04E-02 |
| | ENH | 1.10 | 0.77 | 0.46 | | | 0.92 | 3.02E-134 | 0.00E+00 | | | 2.99E-32 |
| | PRE | 1.13 | ***1.07*** | 0.57 | | | 1.37 | **1.80E-01** | 1.41E-24 | | | 8.26E-04 |
| | ORI | 0.80 | 0.35 | 0.19 | | | 0.49 | 1.26E-10 | 2.51E-18 | | | 5.79E-05 |
| Nipped-B | PRO | 0.96 | 1.06 | 1.08 | 1.08 | 0.67 | 0.88 | 2.85E-11 | 2.16E-27 | 2.16E-12 | 2.94E-240 | 4.13E-30 |
| | High SA PRO | 1.76 | ***1.26*** | ***1.55*** | 1.25 | 1.04 | 1.27 | 1.55E-66 | 6.83E-14 | 3.89E-66 | 3.24E-107 | 7.18E-61 |
| | ENH | 1.49 | 0.89 | 1.31 | 0.88 | 1.71 | 0.88 | 9.42E-253 | 4.25E-30 | 4.50E-247 | 2.82E-36 | 8.51E-269 |
| | PRE | ***1.89*** | ***1.31*** | ***1.57*** | ***1.43*** | ***2.17*** | ***1.66*** | 4.15E-16 | 4.02E-07 | 1.44E-11 | 2.58E-04 | **1.25E-03** |
| | ORI | 0.86 | 0.35 | 0.48 | 0.20 | 0.76 | 0.32 | 1.63E-11 | 1.93E-05 | 4.62E-13 | **3.84E-01** | 1.12E-12 |
| MED30 | PRO | 0.90 | 1.38 | 0.71 | | | 0.79 | 3.12E-234 | 1.03E-52 | | | 3.78E-23 |
| | High SA PRO | 2.15 | 2.87 | 1.99 | | | 1.87 | 5.93E-43 | 3.20E-05 | | | 3.90E-10 |
| | ENH | 2.26 | 2.84 | 2.09 | | | 1.79 | 4.23E-100 | 6.71E-14 | | | 4.38E-63 |
| | PRE | ***2.54*** | ***3.29*** | ***2.23*** | | | ***2.66*** | 3.20E-20 | 1.08E-06 | | | **6.76E-02** |
| | ORI | 0.79 | 0.71 | 0.68 | | | 0.29 | **6.59E-01** | **4.37E-01** | | | 5.47E-06 |
| Fs(1)h | PRO | 1.19 | | | | | 0.49 | | | | | 0.00E+00 |
| | High SA PRO | 1.66 | | | | | ***0.77*** | | | | | 2.17E-183 |
| | ENH | ***1.73*** | | | | | ***0.75*** | | | | | 0.00E+00 |
| | PRE | 1.24 | | | | | ***0.78*** | | | | | 3.11E-23 |
| | ORI | 1.69 | | | | | 0.34 | | | | | 5.16E-20 |

red = UP
blue =DOWN
green = not significant
***bold italics = features with highest median values***

**Supplemental Table S2.** Median values of Rad21, SA, Smc1, Nipped-B, MED30, and Fs(1)h enrichment at promoters, enhancers, PREs and early replication origin centers in control and treated BG3 cells with statistical tests.

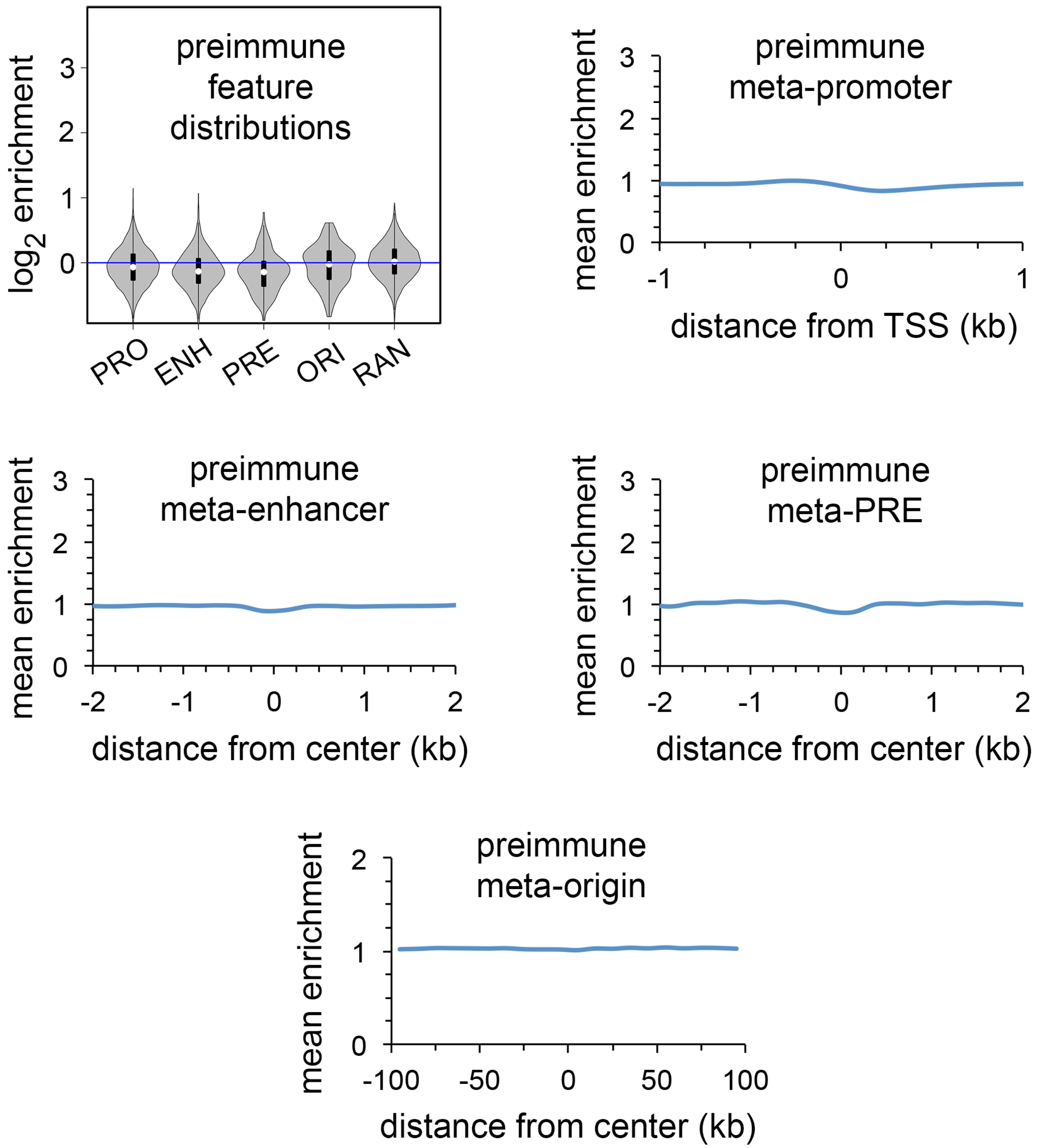


**Supplemental Figure S1.** ChIP-seq with preimmune serum in BG3 cells. Guinea pig preimmune serum ChIP-seq gives no significant enrichment of active promoters (PRO) enhancers (ENH) Polycomb Response Elements (PRE) centers of early DNA replication origins (ORI) or randomly positioned sequences (RAN).

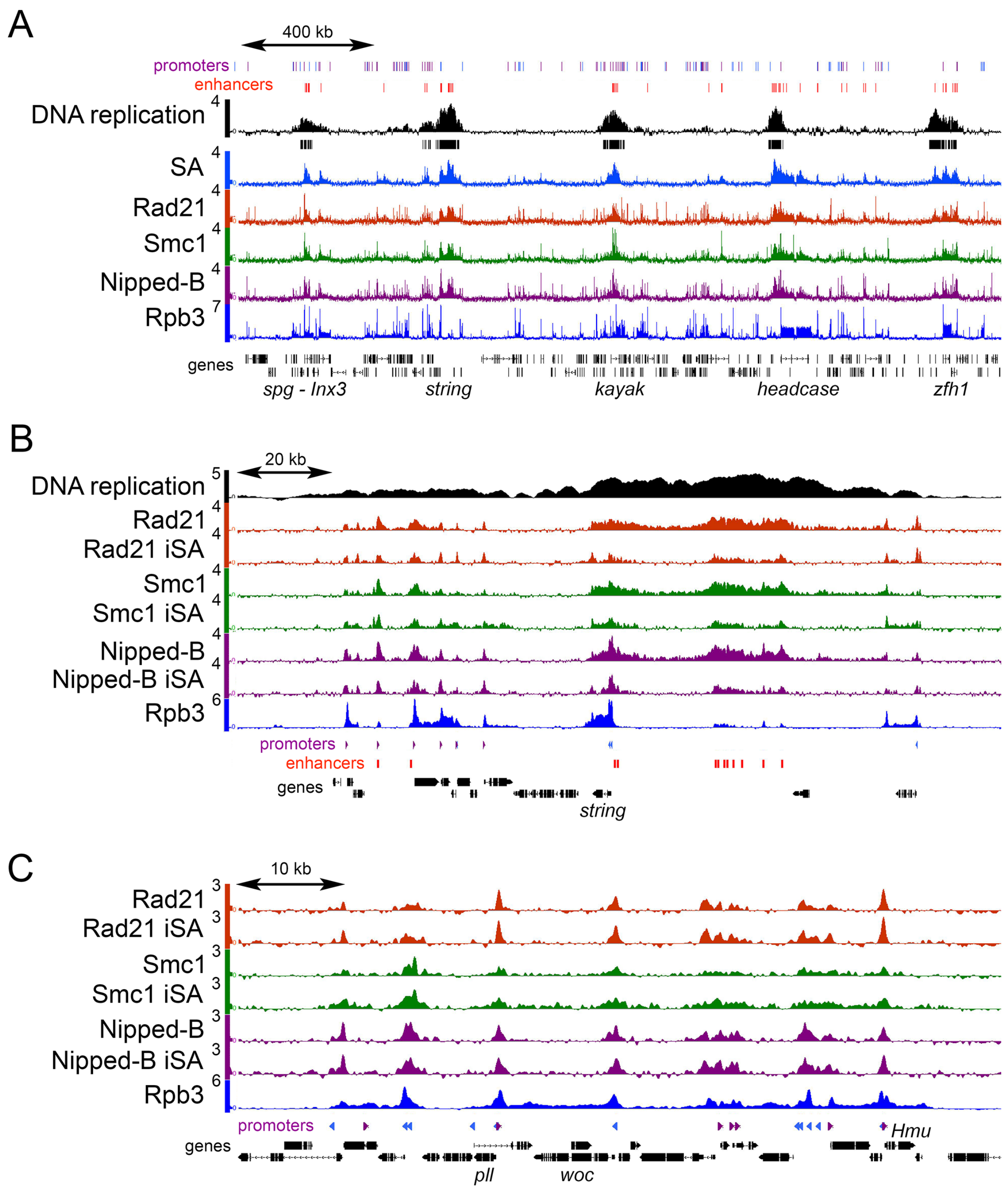


**Supplemental Figure S2.** Early DNA replication origins align with clusters of enhancers and SA depletion selectively reduces Rad21, Smc1 and Nipped-B association with enhancers and origin-proximal promoters in BG3 cells. (*A*) Early DNA replication origins overlapping clusters of enhancers associated with the *spg - Inx3*, *string*, *kayak*, *headcase* and *zfh1* genes. All scales are $log_2$ enrichment and the bars underneath the DNA replication track indicate regions of early DNA synthesis in the $95^{th}$ percentile over regions ≥300 bp in size. (*B*) Rad21, Smc1 and Nipped-B reductions at the *string* enhancers and origin-flanking promoters upon SA depletion (iSA). (*C*) Minor increases in Rad21 and Smc1 at some origin-distal promoters. Enrichment is plotted as $log_2$ values so increases may appear smaller than they are.

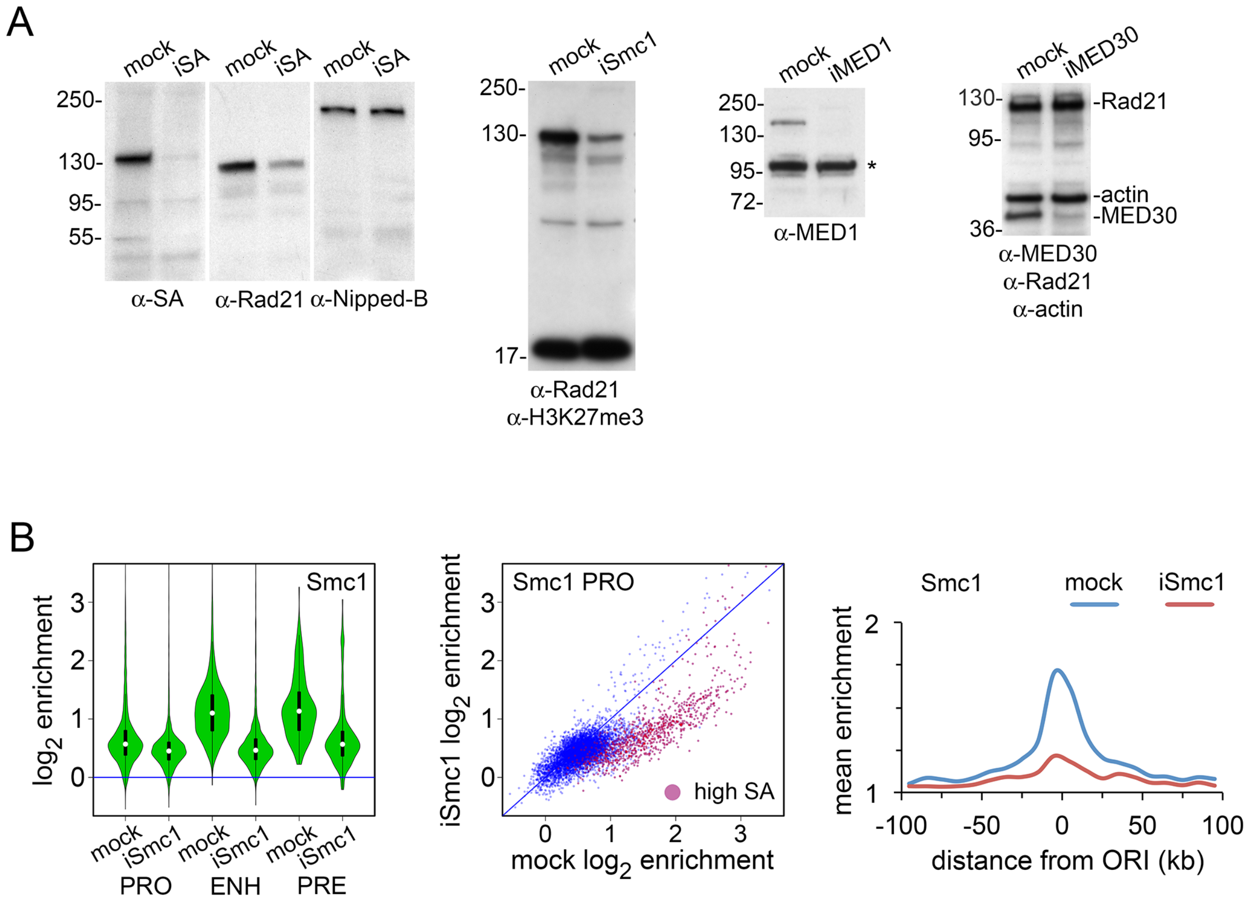


**Supplemental Figure S3.** Effects of RNAi depletions on protein levels or chromosome association in BG3 cells. (*A*) Western blots. Molecular weight markers in kD are indicated left of the panels. The three left panels are duplicate blots of whole cell extracts from mock-treated cells and cells depleted for SA (iSA) probed with the antibodies indicated underneath each panel. The fourth panel from the left is a western blot of whole cell extracts from mock and Smc1-depleted cells (iSmc1) probed with Rad21 and H3K27me3 antibodies. The fifth panel is a western blot of whole cell extracts from mock cells and cells depleted for the MED1 Mediator subunit (iMED1) probed with MED1 antibody. The asterisk (*) indicates a non-specific band recognized by the antiserum. The right panel is a western blot of whole cell extracts of mock-treated cells and cells depleted for MED30 (iMED30) probed simultaneously for MED30, Rad21 and actin. (*B*) Smc1 ChIP-seq of Smc1-depleted (iSmc1) cells. The violin plot distributions, promoter dot plots and meta-analysis of early DNA replication origins show globally reduced Smc1 occupancy leaving similar residual levels at all features. All decreases are statistically significant (Supplemental Table S2). ChIP-seq track examples are in Supplemental Fig. S4.

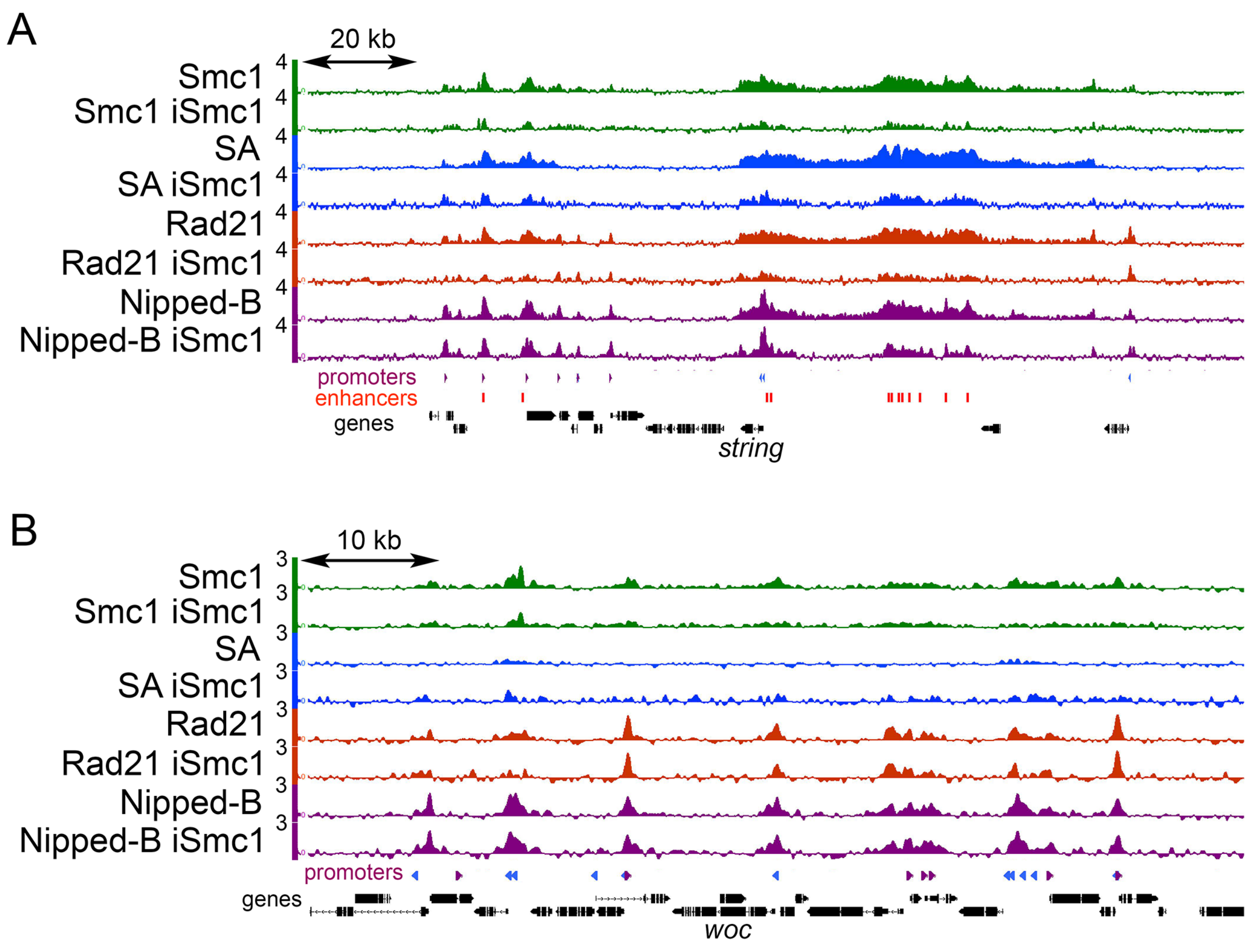


**Supplemental Figure S4.** Smc1 depletion selectively reduces cohesin levels on enhancers and origin-proximal promoters in BG3 cells. (*A*) Effects of Smc1 depletion (iSmc1) on Smc1, SA, Rad21 and Nipped-B occupancy in origin-proximal region containing the *string* gene. (*B*) Effects of Smc1 depletion on Smc1, SA, Rad21, and Nipped-B association in origin-distal regions containing the *woc* gene.

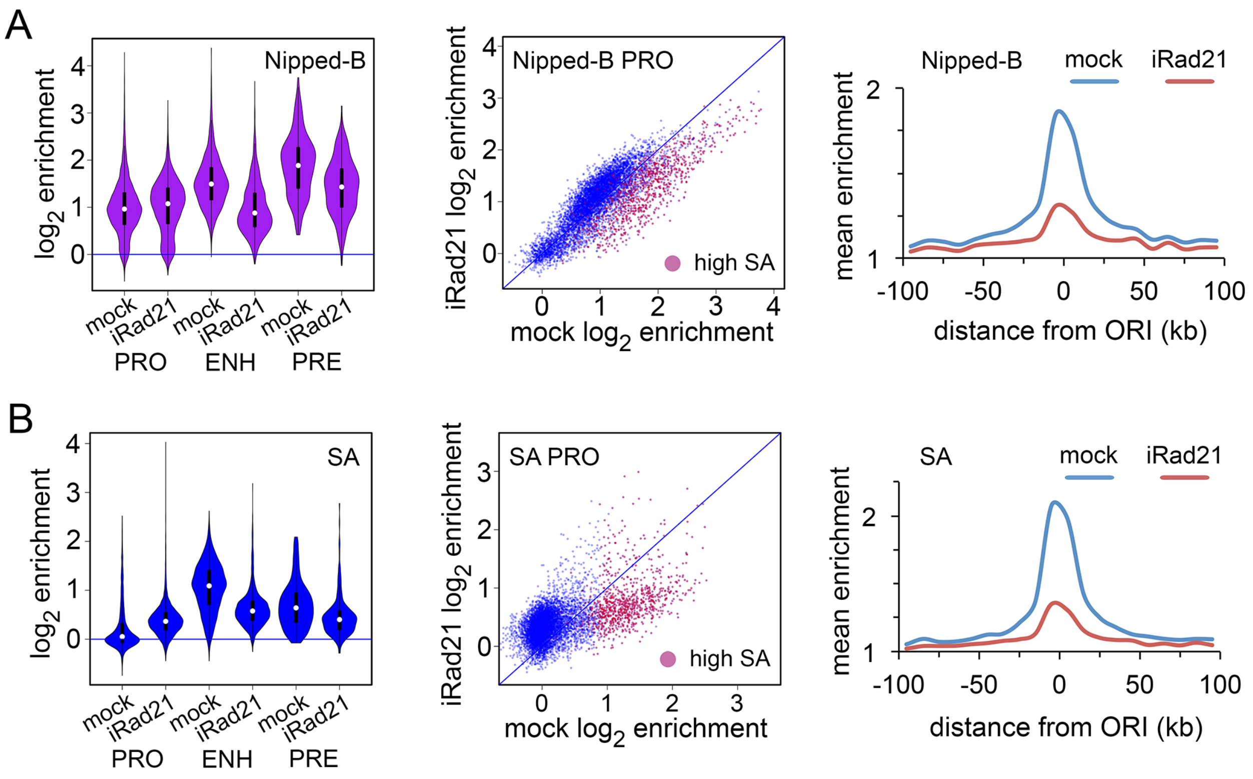


**Supplemental Figure S5.** Rad21 facilitates Nipped-B and SA association with enhancers, high SA promoters and origin-proximal regions in BG3 cells. (*A*) Effects of Rad21 depletion (iRad21) on Nipped-B occupancy. (*B*) Effects of Rad21 depletion on SA occupancy. All changes in Nipped-B and SA occupancy are statistically significant (Supplemental Table S2).

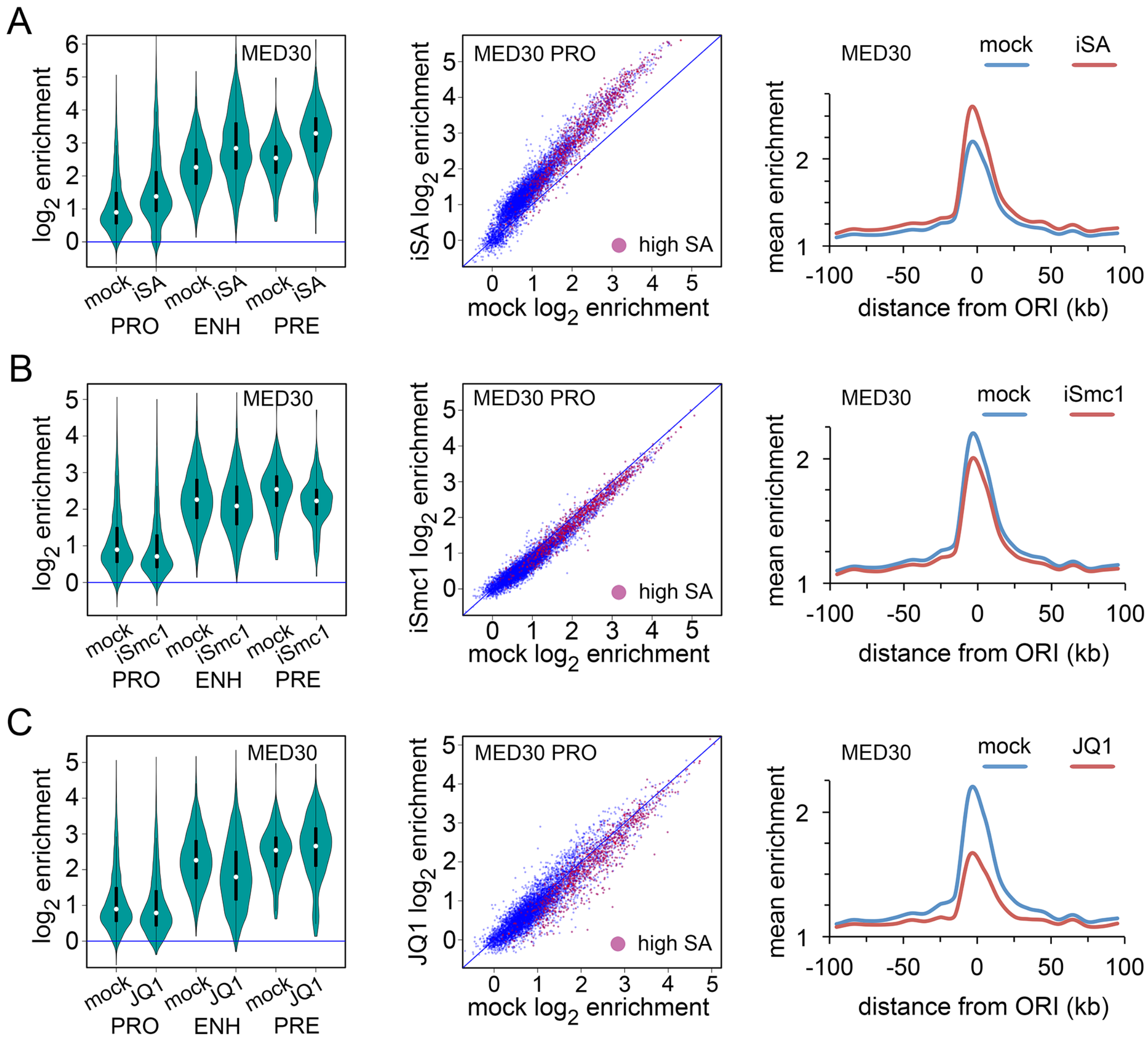


**Supplemental Figure S6.** SA inhibits MED30 association with functional features in BG3 cells. (*A*) Effects of SA depletion (iSA) on MED30 occupancy. (*B*) Effects of Smc1 depletion on MED30 occupancy. (*C*) Effects of JQ1 treatment on MED30 occupancy. All effects on MED30 occupancy are statistically significant except for the effects of SA or Smc1 depletion at the centers of early replication origins, and the effect of JQ1 treatment at PREs (Supplemental Table S2).

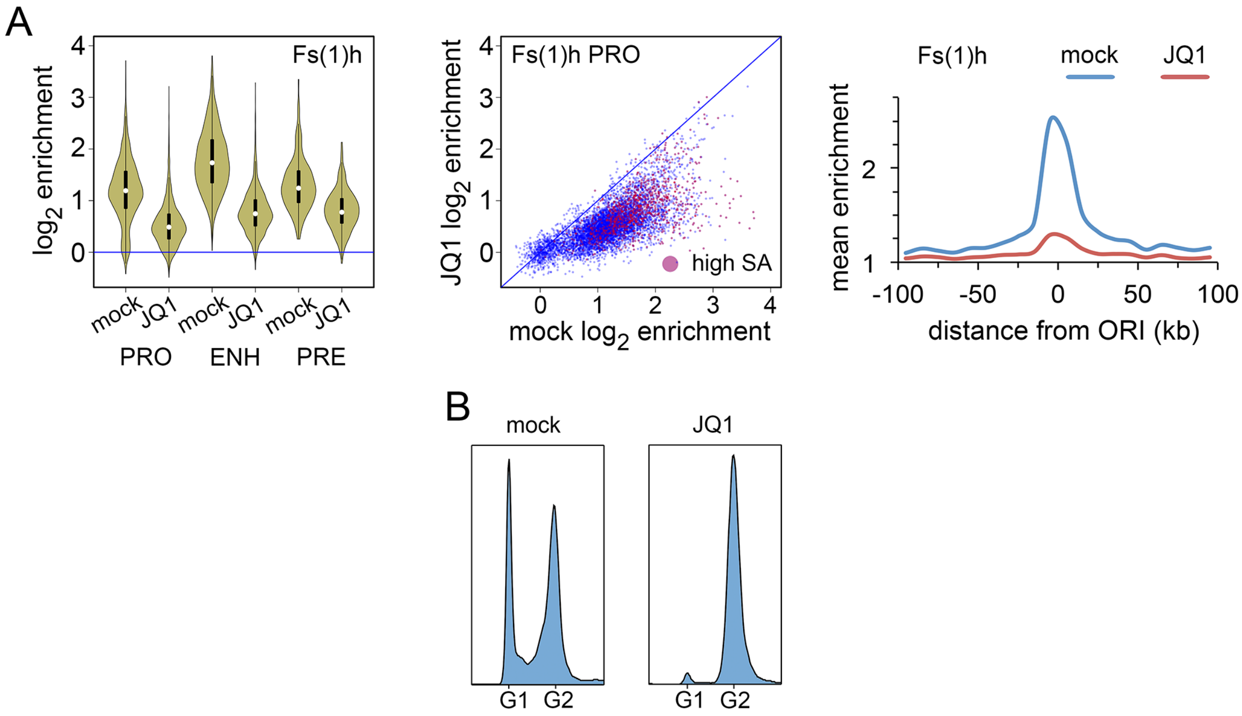


**Supplemental Figure S7.** Effects of JQ1 on Fs(1)h chromosome association and cell cycle in BG3 cells. (*A*) Fs(1)h ChIP-seq after JQ1 treatment (10 micromolar for eight hours). All decreases in Fs(1)h occupancy are statistically significant (Supplemental Table S2). (*B*) Effects of JQ1 treatment on the cell cycle. FACS analysis comparison of mock-treated to JQ1-treated BG3 cells shows that JQ1 arrests cells in G2.
